# Physiological activity within peripheral nerves influences neural output in response to electrical stimulation: an *in vivo* study

**DOI:** 10.1101/2024.12.17.628956

**Authors:** Lauren R. Madden, Richard Liu, Sergiu Ivanescu, Tim M. Bruns

## Abstract

Neuromodulation therapies are often applied to peripheral nerves. These nerves can have physiological activity that interacts with the activity evoked by electrical stimulation, potentially influencing targeted neural output and clinical outcomes. Our goal was to quantify changes in sensory neural unit activity in response to variations in electrical stimulation frequency and amplitude. In a feline model, we applied cutaneous brushing to evoke pudendal nerve afferent activity with and without electrical stimulation via a pudendal nerve cuff electrode. We recorded neural output with microelectrode arrays implanted in ipsilateral sacral dorsal root ganglia (DRG). Combined inter-spike interval distributions for all DRG units showed ranges of flattening, increases, and shifts in response to electrical stimulation. These distributions and changes within them due to electrical stimulation were largely driven by a select few units. Mixed-effects models revealed that quicker firing units generally decreased in firing rate in response to electrical stimulation and, conversely, slower firing units increased in firing rate. A unit’s underlying firing rate also drove the magnitude of change in mean output firing rate in response to stimulation. Further, the models reported a small, negative correlation between the output mean unit firing rate and the applied electrical stimulation frequency. These results demonstrate the potential impact of electrical stimulation on underlying neural firing activity and output. Peripheral neuromodulation may normalize abnormal firing patterns in nerves contributing to pathological disorders or alter unrelated physiological activity in off-target neurons. These factors should be considered when selecting neuromodulation settings in animal subjects and human patients.

## Introduction

Peripheral neuromodulation devices are used as second-or third-line treatments for a wide variety of diseases and disorders. Sacral neuromodulation for bladder dysfunctions, vagus nerve stimulation for epilepsy, and percutaneous peripheral nerve stimulation for chronic pain constitute some of the FDA-approved treatments that employ electrical stimulation of peripheral nerves. There are gaps in understanding of the mechanisms that explain how stimulation of a given peripheral nerve mitigates the disorder a therapy is intended to target. For example, sacral neuromodulation reportedly activates spinal cord reflexes and modulates brain activity that represents bladder-related sensations such as fullness and urge, yet the activation and involvement of direct pelvic floor efferents is currently a topic of discussion (De Wachter et al., 2020). Further, clinical reviews of various peripheral neuromodulation therapies report anywhere from 14 - 41% of patients who experience a loss of effectiveness of their devices over time or who do not experience any benefit after implantation (Gandhi et al., 2021; Helm et al., 2021; Siegel et al., 2018; Toffa et al., 2020).

Electrical stimulation is often administered to nerves that have ongoing physiological activity. This activity may be pathological, non-pathological, or, particularly with functionally mixed nerves, can be off-target and unrelated to the condition to be treated. In either case, peripheral neuromodulation therapies generally aim to modulate the output of neurons that transmit relevant information to the central nervous system or targeted organs or muscles. Vagus nerve stimulation researchers, for instance, have teased out details such as fiber type recruitment, neurotransmitter release, and vagal projection anatomy to elucidate its mechanisms for treatment of epilepsy (Carron et al., 2023). However, the complexity and breadth of vagal connectivity throughout both the brain and the body, and thus the broad scope of information transmission within this nerve, has still presented challenges for device programming and avoiding off-target effects (Groff et al., 2020; Nicolai et al., 2020). How the physiological activity of axons within a nerve is modulated in response to different stimulation settings warrants thorough investigation.

Computational studies have described relationships between a neuron’s endpoint output (the endpoint being, for example, a sensory neuron’s primary synapse) and the ratio of its physiological firing rate to electrical stimulation rate (Crago and Makowski, 2014; Sadashivaiah et al., 2018). These studies highlighted the prevalence of four different interactions that can occur between physiologically induced action potentials (PAPs) and the bidirectional electrical stimulation induced action potentials (ESAPs) within a physiologically active and electrically stimulated axon (*Figure 1*). The first is collision block, during which an antidromic ESAP collides with a PAP approaching the stimulation site along the axon, and they annihilate each other. This results in only the orthodromic ESAP reaching the fiber’s endpoint. The second is resetting by stimulation, during which the antidromic ESAP reaches the source of the PAP, such as a neural sensory ending, and prevents a PAP from being generated. This interaction also results in only the orthodromic ESAP reaching the endpoint. The third type of interaction is refractory block, during which the hyperpolarized phase of the PAP is passing the site of stimulation during a stimulating pulse, preventing both ESAPs from occurring due to the axon being in a refractory state. In this case, only the PAP reaches the endpoint. The fourth interaction, summation, occurs when the antidromic ESAP does not collide with a PAP before reaching the neural source, nor does it prevent a PAP from being generated. This leads to the orthodromic ESAP and both the preceding and following PAPs reaching the endpoint.

**Figure 1:**
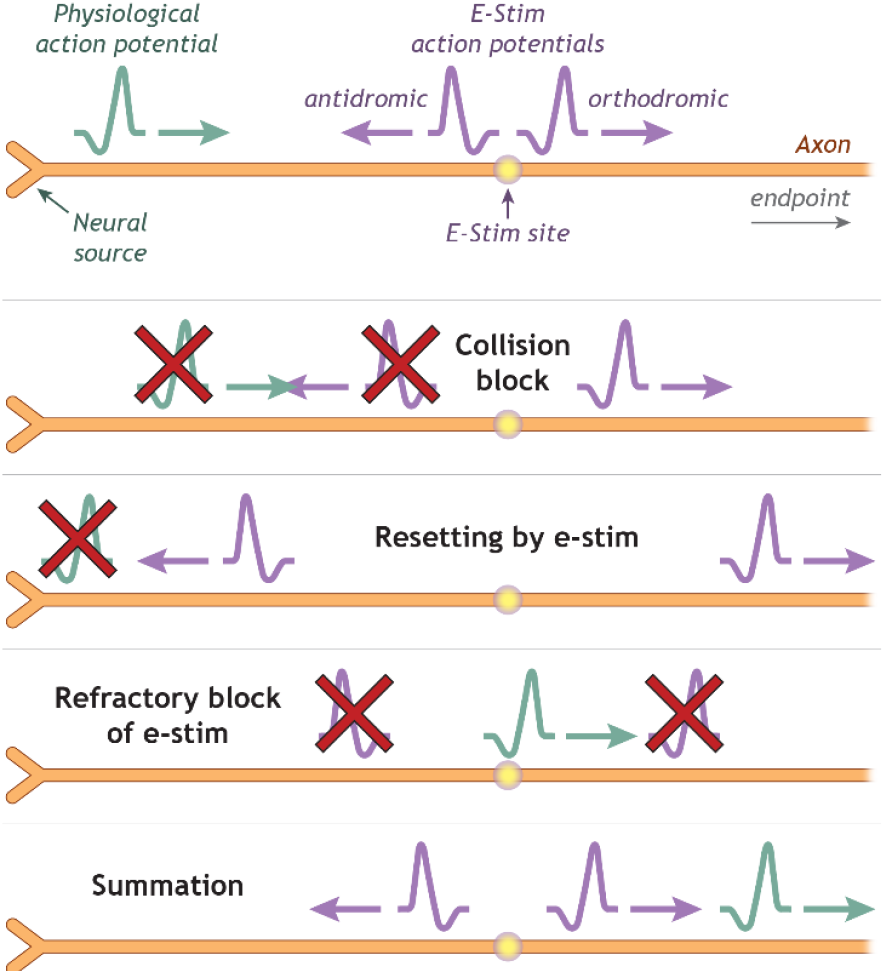
A schematic showing the four different types of interactions between a PAP (green) and ESAPs (purple). The orange axon represents a sensory axon, where the sensory ending is depicted on its left.

The computational studies by Crago and Makowski 2014 and Sadashivaiah et al., 2018 provide a contextual framework for how the output of individual physiologically active axons would be expected to change while undergoing electrical stimulation. However, these unitary relationships have yet to be investigated *in vivo*. Here, we quantified alterations in firing output of physiologically active neural units in response to various stimulation frequencies and two stimulation amplitudes in a feline model. We applied cutaneous brushing to evoke PAPs, applied electrical stimulation with a cuff electrode to an apposite nerve to evoke ESAPs, and recorded neural unit endpoint output in the respective dorsal root ganglia (DRG) with microelectrode arrays. We compared the inter-spike-interval (ISI) distributions and changes in firing rate of neural units recorded during brushing while applying stimulation to the distributions recorded during brushing without stimulation in order to quantify the modulation of the active units’ output. Our goal was to provide experimental context to describe the heterogeneous response patterns of physiologically active fibers undergoing different peripheral nerve stimulation frequencies and amplitudes.

## Methods

### Animals

All procedures were approved by the University of Michigan Institutional Animal Care and Use Committee (IACUC), in accordance with the National Institute of Heath’s guidelines for the care and use of laboratory animals. We collected data across five experimental sessions using three adult, domestic, purpose-bred short-hair male cats (1.3 ± 0.3 years old, 4.9 ± 0.6 kg). Animals were free-range housed with up to 6 other cats in a 38 m^2^ room with controlled temperature (19-21 °C) and relative humidity (35-60%), food and water available ad libitum, and a 12-hour light/dark cycle. Staff interaction and toys provided enrichment for the animals.

### Surgical procedure

Animals were initially anesthetized with ketamine (6.6 mg/kg), butorphanol (0.66 mg/kg), and dexmedetomidine (0.011 mg/kg) via a mixture injected intramuscularly. They were then intubated, laid prone, and maintained on isoflurane anesthesia (0.5-2%) during surgical procedures and data collection sessions. We continuously monitored temperature, heart rate, respiratory rate, end-tidal CO_2_, O_2_ perfusion, and intra-arterial blood pressure using a SurgiVet vitals monitor (Smiths Medical, Dublin, OH). Throughout the procedure, fluids (1:1 ratio of lactated Ringer’s solution and 5% dextrose) were infused intravenously via the cephalic vein at a rate of 10 ml/h. We inserted a 3.5 Fr diameter catheter into the bladder through the urethra for fluid draining. We made a postero-lateral incision to expose the pudendal nerve, on which we placed a bipolar stimulation cuff fabricated in our lab (2-mm inner diameter, 2-2.5 mm wire separation). We made a midline dorsal incision to expose the L7-S3 vertebrae and performed a partial laminectomy to access the ipsilateral S1 and S2 DRG. We implanted either one or two 32-channel Utah microelectrode arrays (Blackrock Neurotech, Salt Lake City, UT), depending on the availability of devices on-hand, to either or both DRG with a pneumatic inserter (Blackrock Neurotech, Salt Lake City, UT). We recorded neural signals with the microelectrode arrays along with a Grapevine neural interface processor and Trellis Acquisition Software (Ripple Neuro, Salt Lake City, UT) at a rate of 30 kHz. All microelectrode array channels from which we sorted neural units had impedances below 200 kΩ.

Each data collection session took place while the animal was under isoflurane anesthesia during either one of two procedural scenarios (*Table S1*). One scenario was during a device implant procedure after which the animal recovered from the surgery, as previously described (Khurram et al., 2017), prior to testing for separate experimental goals. The second type of scenario was a terminal procedure, after which the animal was euthanized with a sodium-pentobarbital solution (36 mg/kg, intravenously). In both scenarios, any microelectrode arrays used for collecting data were newly implanted. Test sessions between an implant procedure and a terminal procedure were not possible.

### Data collection

To evoke PAPs, we used a cotton swab fixed to a high-resolution linear motor actuator (model M-235.5DD, Physik Instrumente, Auburn, MA) that was mounted to an optical board and aligned to brush either the perineal region, the scrotum, or the anus. The actuator was controlled by a MATLAB (MathWorks, Inc., Natick, MA) script to apply cutaneous brushing at regular 2 or 4 second intervals, where the actuator moved back and forth across 10 mm or 20 mm distances, respectively, at a velocity of up to 15 mm/s. We recorded unit extracellular voltage waveforms via the microelectrode arrays implanted in the ipsilateral DRG for 40-60 second trials captured with the Ripple Grapevine neural interface processor and Ripple Trellis acquisition software. The Trellis software applied a band-pass filter (250 Hz to 7.5 kHz) to the raw data prior to automated detection of spike waveforms that exceeded a threshold of 3.5 times the root mean square of the voltage on each microelectrode channel. See *Figure 2* for an experimental setup diagram. For each respective trial set, we recorded at least one trial where only cutaneous brushing was applied without electrical stimulation to obtain a physiological baseline distribution of PAP ISIs.

**Figure 2:**
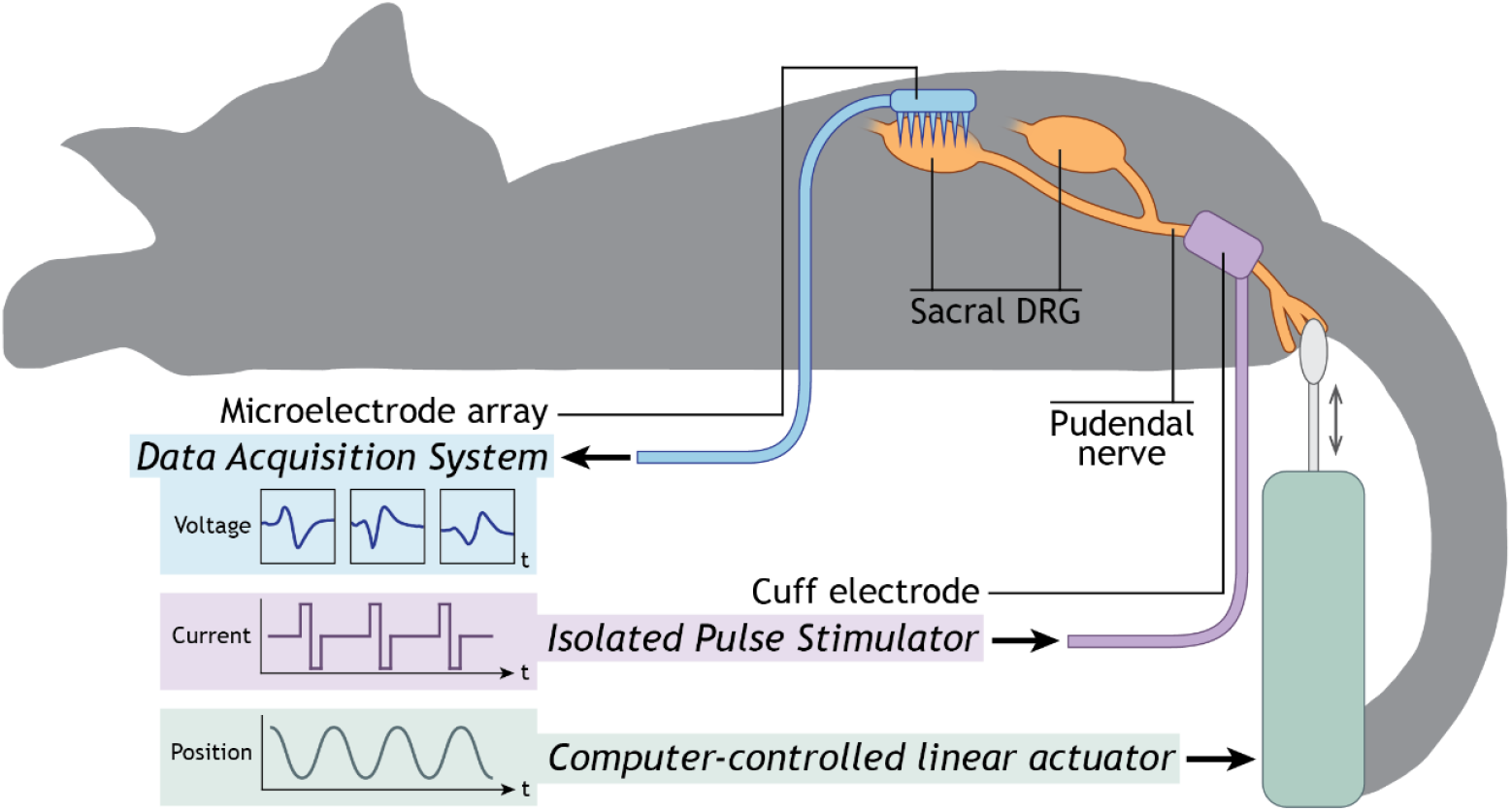
Experimental setup diagram. PAPs were evoked via cutaneous brushing with a computer-controlled linear actuator (green). ESAPs were evoked via a cuff electrode connected to an isolated pulse stimulator (purple). Endpoint action potentials were recorded with the Ripple Grapevine/Trellis system connected to a Utah array (blue) implanted in sacral dorsal root ganglia. Sacral DRG and pudendal nerve are shown in orange. Not to scale.

To evoke ESAPs, we applied functionally equivalent 200 or 210 µs biphasic electrical pulses with the bipolar cuff electrode around the pudendal nerve, which was controlled by an isolated pulse stimulator (model 2100, A-M Systems, Carlsborg, WA). For each respective data collection session, we first identified the motor activation threshold amplitude by slowly ramping up the isolated pulse stimulator amplitude from zero until we visually observed activation of the external anal sphincter (EAS). We used the minimum amplitude required to elicit EAS activation, i.e., the motor activation threshold amplitude (MT), along with twice the motor activation threshold amplitude (2xMT) during trials that administered electrical stimulation. We selected logarithmically spaced stimulation frequencies within 0.5 - 40 Hz for stimulation trials to increase the effect resolution of neural output during lower stimulation frequency values.

A summary of parameter information across animals, sessions, and trial sets can be found in *Table S1*. For the single data collection session with Animal 1, we recorded 10 total electrical stimulation trials with five stimulation frequencies spaced from 2 - 40 Hz and two stimulation amplitudes, MT and 2xMT, where each combination of frequency and amplitude was used for one trial. We repeated this across three total trial sets where we brushed the anus, scrotum, and perineal region during each set, respectively. For the data collection sessions with Animals 2 and 3, we applied 10 stimulation frequencies spaced from 0.5 - 30 Hz and two stimulation amplitudes, MT and 2xMT. We recorded 20 total trials for each trial set, where each combination of stimulation frequency and amplitude was used for one trial. For the first data collection session with Animal 2, we recorded two sets of trials, where we applied cutaneous brushing to the perineal region and anus for each trial set, respectively. For the second data collection session with Animal 2, we applied brushing to the scrotal region only. During each data collection session with Animal 3, we only collected a single trial set while brushing the perineal region.

We collected all trial recordings for a given data collection session within one to three hours. Because we implanted the recording microelectrode array at the beginning of each data collection session, all neural units recorded during a given session were assumed to be unique from units collected during the alternative session with the same animal. We performed a post-mortem dissection on Animal 3 to measure the distance between the cuff electrode and DRG, which was 5.6 cm for S1 and 5.1 cm for S2. We used these distances for calculating the conduction velocity for units that responded directly to electrical stimulation in each animal.

### Neural data analysis

We used Offline Sorter (Plexon, Dallas, TX) to manually sort neural spike waveforms within individual electrode channels into respective units (Khurram et al., 2017). Manual sorting enabled discrimination between neural unit spikes, noise, and electrical stimulation artifacts via visual inspection of their waveforms (shape and amplitude) and timing (e.g. ambient noise artifacts were identified with a frequency of 60 Hz and harmonics of it). We used neural unit spike shape, assisted by principle component analysis, to sort neural waveforms recorded on the same channel. We maintained intra-channel unit labels across all trials within a data collection session. We exported the sorted data to obtain unit spike timestamps for each trial, which we then used to calculate ISIs in MATLAB. ISIs longer than 1 second were omitted for each unit to eliminate the amount of time between units’ responses to consecutive brushing events during a trial. We used MATLAB’s kernel density estimation function to generate probability density functions (PDFs) for each unit’s ISI distribution during a given trial.

We calculated the mean conduction period based on the time intervals between each electrical stimulation artifact and the subsequent spike waveform for each stimulation-responding unit during its lowest stimulation frequency trial at 2xMT amplitude, for five stimulation events while brushing was not occurring. The mean conduction period for each unit was divided by the estimated distance between its respective DRG and the cuff electrode, as based on the measures obtained from Animal 3, to obtain the approximate mean conduction velocity for a given unit. For some units it was unclear if they responded directly to electrical stimulation due to continuous firing behavior, so these units were omitted from further analysis.

We use *T*_*n*_ to denote a distribution of physiological ISIs (i.e., intervals between PAPs induced by brushing without electrical stimulation) and *T*_*e*_ to denote a distribution of endpoint ISIs collected during any trial. Alternatively, we use *R*_*n*_ to denote a distribution of physiological instantaneous firing rates (i.e., reciprocals of *T*_*n*_) and *R*_*e*_ to denote a distribution of endpoint instantaneous firing rates (i.e. reciprocals of *T*_*e*_). Note that for trials where only brushing was applied without electrical stimulation, *R*_*e*_ equals *R*_*n*_ (and *T*_*e*_ equals *T*_*n*_), as we used *R*_*e*_ values from the brushing only trials to determine the underlying firing rate distribution (*R*_*n*_). We use *R*_*s*_ to denote the applied electrical stimulation frequency during a given trial and, correspondingly, *T*_*s*_ to denote the inter-pulse period of an applied electrical stimulation frequency, where *T*_*s*_ equals 1/*R*_*s*_.

### Statistical Analysis

To determine the mean firing rate (or mean ISI) of a unit during a trial, we calculated the log-average of the unit’s instantaneous firing rate (or ISI) distribution to obtain the geometric mean. The geometric mean is a more appropriate representation of the central tendency of distributions that are heavily skewed, as firing rate distributions of neurons often are (Buzsáki and Mizuseki, 2014). The skewness values of individual units’ ISI distribution were determined using MATLAB’s *skewness* function. For a given ISI distribution, we applied a natural logarithm transformation to the distribution and calculated the arithmetic mean of the log-transformed values. We then applied an exponential transformation of the resulting value with Euler’s number to obtain the geometric mean of the ISI distribution. To calculate the mean instantaneous firing rate, we took the reciprocal of each ISI value before performing the log transformation. This calculation is shown in *Equation 1*:

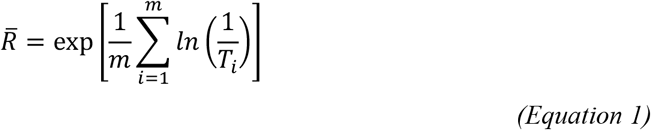

where 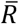 is the geometric mean of instantaneous firing rates (either *R*_*n*_ or *R*_*e*_) from a unit during a given trial, *T*_*i*_ is an ISI value from that unit’s response during the trial (either *T*_*n*_ or *T*_*e*_), and *m* is the number of ISI values from the unit during the trial. Note that *R*_*i*_ = 1/*T*_*i*_, where *R*_*i*_ is an instantaneous firing rate datapoint, and 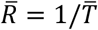, where 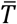 is the geometric mean of ISIs from a unit during a given trial.

We performed right- and two-tailed Student’s t-tests in MATLAB to determine the significance and direction of how given electrical stimulation settings impact specific intervals in a unit’s ISI distribution. For a given t-test, the height of the ISI bin (in terms of normalized probability) in question was calculated for each unit during the brushing only trials. These values were subtracted respectively from each unit’s ISI bin height during a stimulation trial using the settings in question. The null hypothesis for each test was that the mean difference across all stimulation-responding units is zero. We report the test statistics and *p*-values against an α significance level of 0.05 for each test.

Additionally, we used MATLAB to fit the mean firing rate of individual units during given stimulation conditions to two log-linear mixed-effects models. For these models, we only used the data from units that were both modulated by brushing and responded to electrical stimulation. Because of the skewed nature of the response distributions of the data, we fit the linear mixed-effects models to log-transform values of firing rates and electrical stimulation frequencies. The first model used the mean of the log-transformed *R*_*e*_ as the response value. The second model used the difference in the mean of the log-transformed *R*_*e*_ from the mean of the log-transformed *R*_*n*_ (which is equivalent to the log ratio of these means) as the response value. The specification formulas for these two models are written respectively in *Equation 2* and *Equation 3*:

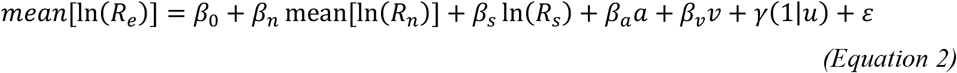

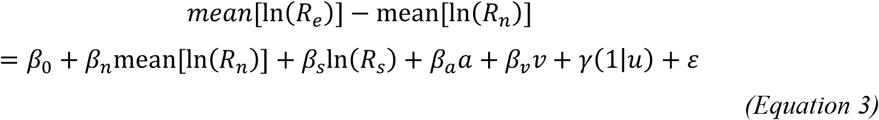

where *a* represents the motor activation threshold amplitude multiplier (valued at either 1 for MT or 2 for 2xMT) used during a given trial, *v* represents the conduction velocity of a unit, *u* represents a unit’s identifier which was treated as a random-effect grouping variable with variance *γ, β*_0_ is the global intercept, *β*_*x*_ is a fixed effect coefficient, and *ε* is the residual variance. Note that an exponential transformation of the response value for *Equation 2* using Euler’s number gives the geometric mean of *R*_*e*_. An equivalent transformation of the response value for *Equation 3* gives the mean of *R*_*e*_ during a trial as a percentage of a unit’s 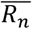 (geometric mean of *R*_*n*_), where a value below 1 (or 100%) indicates a decrease in mean firing rate, and a value above 1 indicates an increase in mean firing rate. We calculated the effect sizes for each coefficient for each model by first normalizing the predictor variable data before compiling it into a table on which we ran MATLAB’s linear mixed-effects model function.

## Results

In total, we sorted data from 84 neural units from all three animals, where 61 responded to brushing and 8 responded to breathing motion (which were likely cutaneous afferents transmitting information from a dermatome that moved with the animal’s body during breathing). We determined a neural unit responded to brushing by whether it changed its firing rate during brushing events. Eighteen of the sorted units did not respond directly to electrical stimulation, and for 28 of the sorted units it was unclear if they responded directly to electrical stimulation due to continuous firing behavior. Thus, we were able to calculate conduction velocities for the remaining 38 units (mean ± standard deviation, 2.11 ± 1.40 m/s; range, 0.35 – 8.0 m/s). Of these, we identified 33 units that were both modulated by brushing and were activated by at least the 2xMT stimulation amplitude (allowing estimation of their conduction velocities). This subset of units was used in subsequent analyses.

Individual unit physiological ISI distributions (i.e. *T*_*n*_, distributions from trials where only brushing was applied, without electrical stimulation) of these 33 units ranged in shape, but virtually all were right skewed (*Figure S1*). Only one unit’s *T*_*n*_ had a skewness of less than 0.5 (3% of the 33 units), and only three units’ *T*_*n*_s had a skewness of less than 1 (9% of the 33 units). The geometric means, medians, and standard deviations of each individual unit’s physiological instantaneous firing rate distribution (*R*_*n*_), along with the conduction velocities of units that responded directly to stimulation, can be found in *Figure 3*. Raster plots and ISI distributions for three example units can be found in *Figure 4*, where each unit’s response is shown for a trial where only brushing was applied, a trial where a 2 Hz stimulation frequency at MT amplitude was applied, and a trial where a 2 Hz stimulation frequency at 2xMT amplitude was applied.

**Figure 3:**
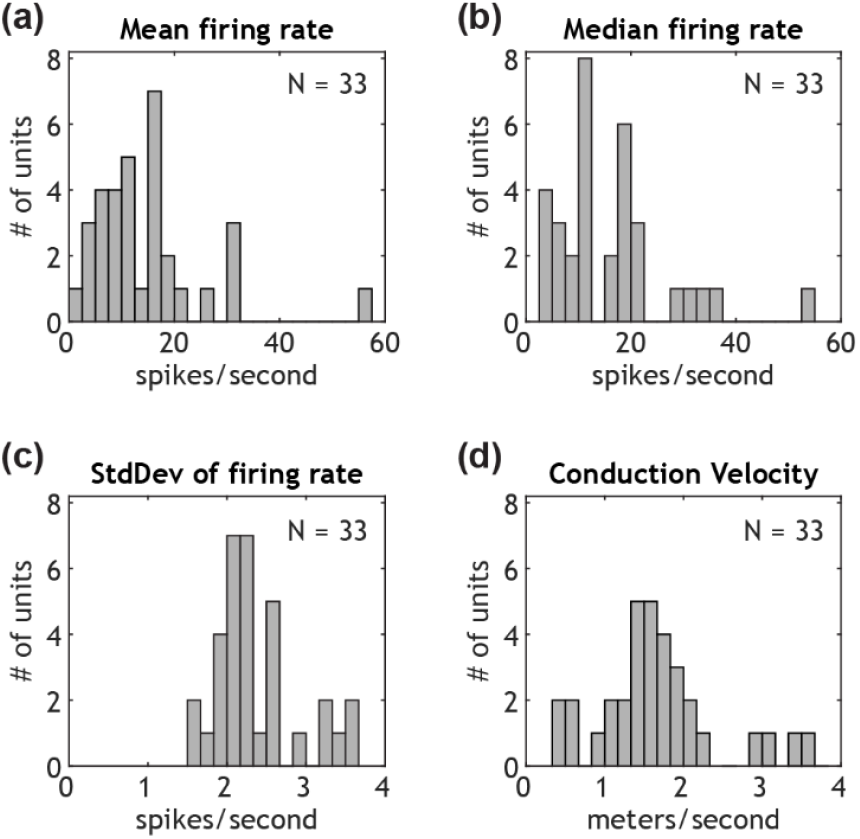
Summary of sorted units that were analyzed for this study. The mean, median, and standard deviation of each unit’s physiological firing rate were calulated using log transformed distributions.

**Figure 4:**
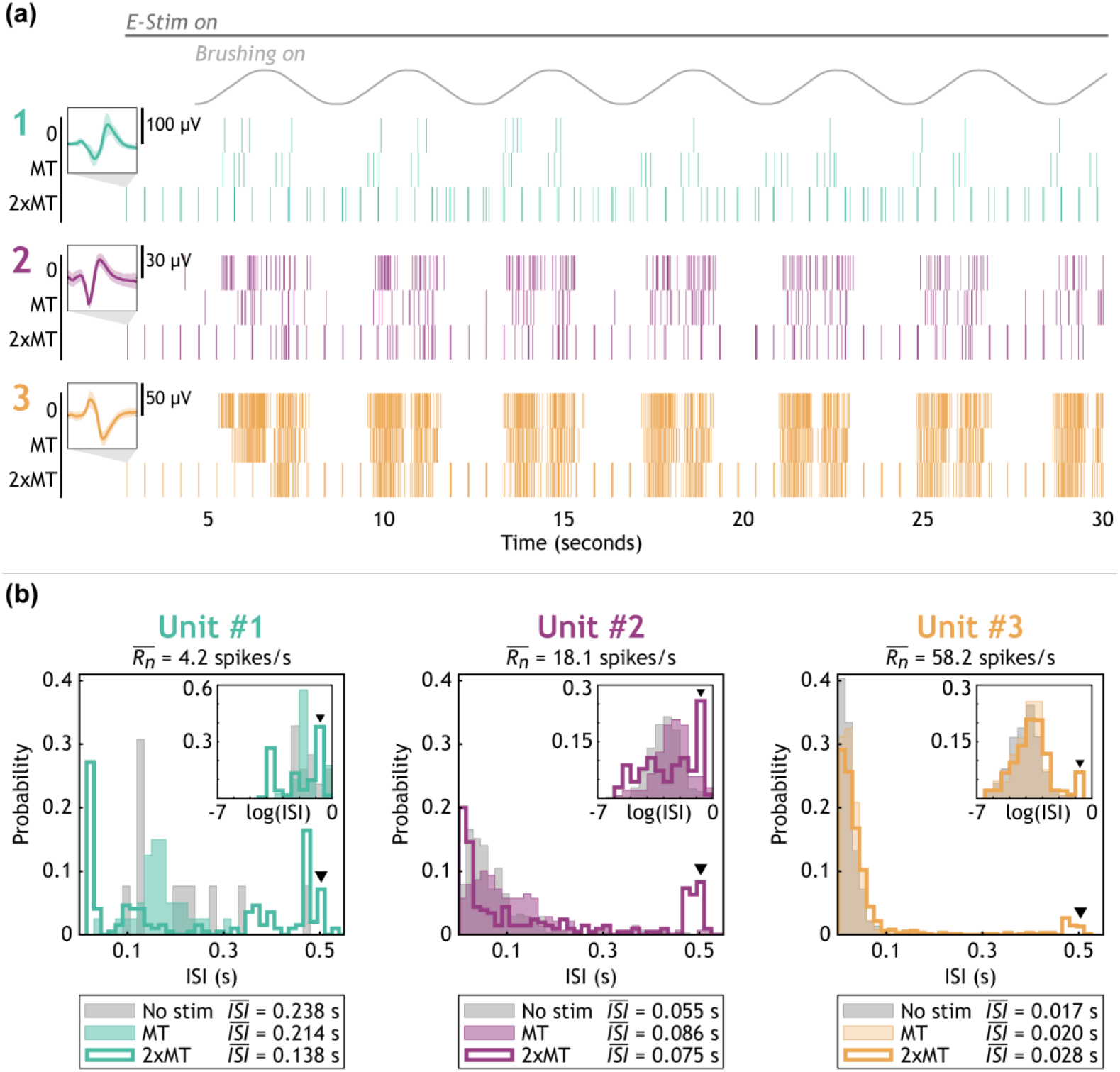
Raster plots (a) and ISI histograms (b) for three example units: one slower firing unit (Unit #1), one unit with a medium firing rate (Unit #2), and one quickly firing unit (Unit #3). Unit waveform insets in (a) have a time scale of 1.7 ms. Trials labeled as “0” or “No stim” are brushing only trials. Data shown from trials where electrical stimulation was applied used a 2 Hz stimulation frequency and are labeled as “MT” for a motor threshold amplitude trial or “2xMT” for a twice motor threshold amplitude trial. The black triangles in (b) indicate the bin that contains T_s_ (0.5 seconds for 2 Hz).

### Collective neural unit responses

We observed several types of response patterns in the ISI distributions combined from all 33 brushing-modulated neural units. These combined distributions during brushing only, 2 Hz, and 30 Hz trials are shown in *Figure 5. Table 1* shows the results of t-tests on unit ISI bin heights respective to this figure. An electrical stimulation frequency of 2 Hz (*Figure 5b* and *5d*) led to a significant increase in very short ISIs. Specifically, the probability of ISIs ranging from 0 to 0.0101 seconds occurring was significantly greater across units during trials applying 2 Hz of electrical stimulation than during brushing only trials (one-sample, right-tailed t-tests for MT trials: t-statistic, 2.67; *p*-value, 0.0059; for 2xMT trials: t-statistic, 2.63; *p*-value, 0.0066). This effect was likely due to summation (*Figure 1*). Correspondingly, the middle range of the distribution (about 0.02 to 0.2 s) reduced with respect to that of brushing only trials. This is illustrated in *Table 1* by the 0.033s and 0.066s unit ISI bin heights during 2 Hz, 2xMT stimulation being significantly lower across units than respective bin heights from the brushing only trials.

**Table 1:** Statistical test outcomes showing impact of electrical stimulation on endpoint ISI bin height in Figure 5.

|  | **2 Hz electrical stimulation** |  | **30 Hz electrical stimulation** |  |
| --- | --- | --- | --- | --- |
| *Amplitude* | *MT* | *2xMT* | *MT* | *2xMT* |
| 0.5s ISI bin<br>(2 Hz $T_s$) | t-stat: 3.1924<br>*p*-value: **0.0032** | t-stat: 3.5385<br>*p*-value: **0.0013** | t-stat: 2.8455<br>*p*-value: **0.0077** | t-stat: 2.1514<br>*p*-value: **0.0391** |
| 1.0s ISI bin<br>(2 Hz $T_s \times 2$) | t-stat: 1.2887<br>*p*-value: 0.2067 | t-stat: -0.1002<br>*p*-value: 0.9208 | t-stat: -0.9164<br>*p*-value: 0.3663 | t-stat: -0.9171<br>*p*-value: 0.3659 |
| 0.033s ISI bin<br>(30 Hz $T_s$) | t-stat: -0.8035<br>*p*-value: 0.4276 | t-stat: -3.2030<br>*p*-value: **0.0031** | t-stat: -0.1311<br>*p*-value: 0.8965 | t-stat: -1.0573<br>*p*-value: 0.2983 |
| 0.066s ISI bin<br>(30 Hz $T_s \times 2$) | t-stat: -0.9207<br>*p*-value: 0.3641 | t-stat: -2.4480<br>*p*-value: **0.0200** | t-stat: 0.2657<br>*p*-value: 0.7922 | t-stat: 1.7059<br>*p*-value: 0.0977 |

**Figure 5:**
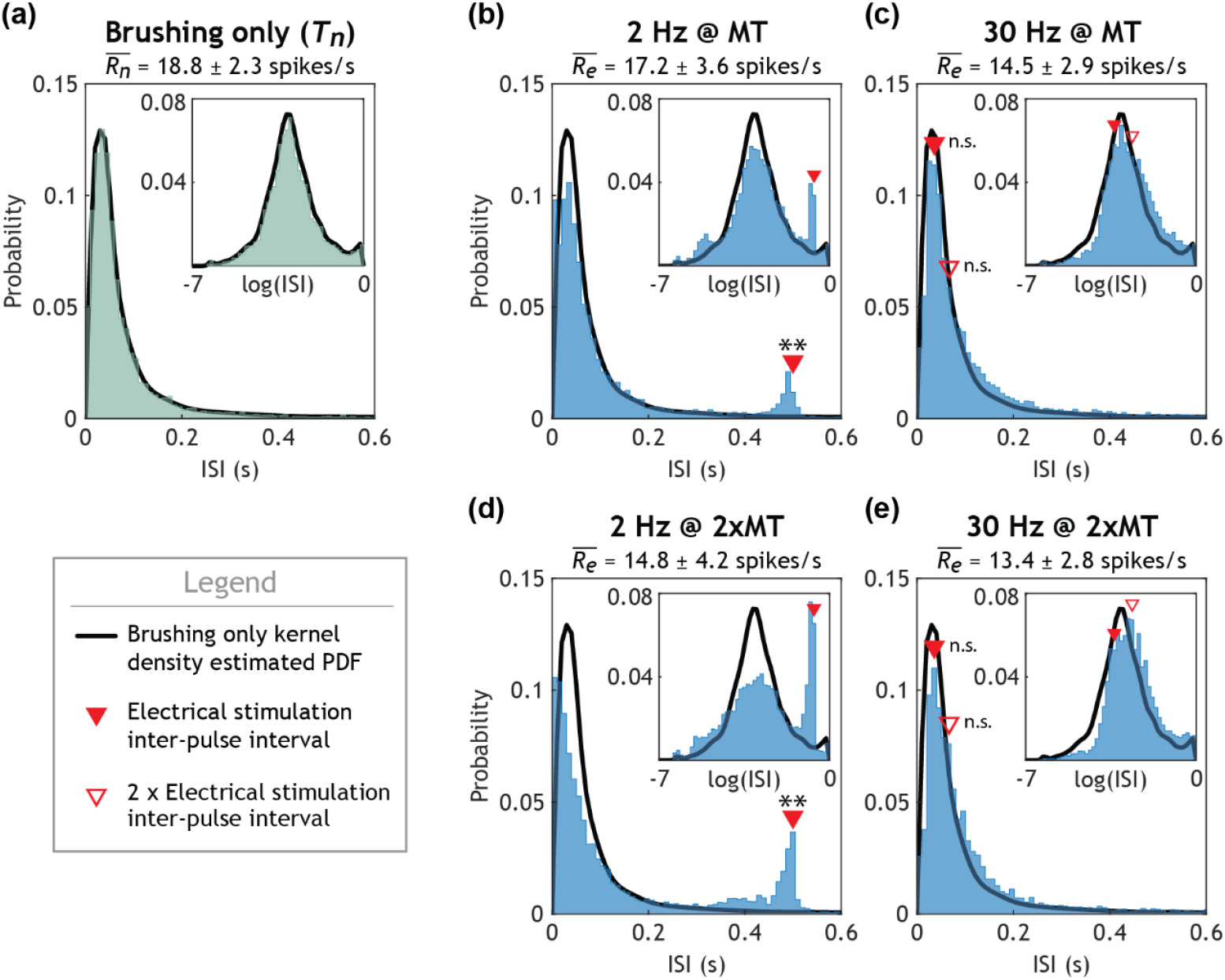
Histograms of ISIs for all brushing modulated units across all animal trials that responded to electrical stimulation. Plot titles show applied electrical stimulation settings (a: brushing only, or b-e: frequency @ amplitude) and the mean ± standard deviation for a given plot’s *R_e_*. The subplot insets show the log transformed ISI distributions for each stimulation condition. The thick black line on all histogram plots indicates the kernel density estimated probability density function (PDF) for the brushing only (physiological) ISI distribution. Solid red triangles indicate the bin that contains the inter-pulse interval between electrical stimulation pulses. Unfilled red triangles indicate the bin that contains twice the inter-pulse interval between electrical stimulation pulses. Asteriks indicate statistical significance of the inter-pulse interval bin height (for a given stimulation condition) being different from that of the brushing only bin height (* p-value < 0.05; ** p-value < 0.01).

Across the middle range of electrical stimulation frequencies we tested, there were added peaks in the ISI distributions that aligned with *T*_*s*_ and multiples of *T*_*s*_ (see *Figure 5b* and *5d, Figures S2* stimulation frequencies 2 Hz – 12 Hz, *Figure S3* stimulation frequencies 2 Hz – 21 Hz, and the 0.5s ISI bin row of the 2 Hz electrical stimulation columns in *Table 1*). It should be noted that while the most prominent added peaks were generally those that match *T*_*s*_, there were small but consistent additional peaks that aligned with multiples of *T*_*s*_. Further, these increased ISI bin regions offset the density of the *T*_*e*_ probability distributions. The most dense region of the combined *T*_*n*_ distribution reduced in the combined *T*_*e*_ distributions during these trials (*Figures S2 and S3*).

Interestingly, in response to the higher stimulation frequencies tested, we observed a small but notable reduction of shorter (< 0.1s) ISI values within the distributions in conjunction with a shift of the entire distributions (see *Figures 5c* and *5e, Figure S2* stimulation frequencies 19 and 30 Hz, and *Figure S3* stimulation frequencies 30 and 40 Hz). The ISI distributions for these same stimulation frequencies also displayed an increase in bin height for ISIs above 0.1s (*Figures 5c* and *5e, Figure S2* stimulation frequencies 19 and 30 Hz, and *Figure S3* stimulation frequencies 30 and 40 Hz). This is illustrated in *Table 1*, where the 0.5s ISI bin height for 30 Hz stimulation trials significantly increased for both MT and 2xMT amplitudes.

### Contribution of individual units to overall endpoint output distributions

We used Pareto charts to quantify the number and degree to which individual units contributed to the combined ISI distributions for each electrical stimulation setting. The combined endpoint output ISI distributions of brushing-modulated units for a given data collection session were often dominated by a handful of units across stimulation settings (*Figure 6*). Seven of the 12 units (58.3%) from the data collection session with Animal 1 and five of the 19 units (26.3%) from the first data collection session with Animal 2 that were both modulated by brushing and activated by electrical stimulation were within the top 10 contributors to the overall combined ISI distribution across all stimulation conditions, including brushing only trials. Percentages of brushing- and stimulation-modulated units for individual data collection sessions with Animal 1 and Animal 2 that were consistently either top or bottom contributors to the respective data collection sessions’ ISI distributions are shown in *Table S2*.

**Figure 6:**
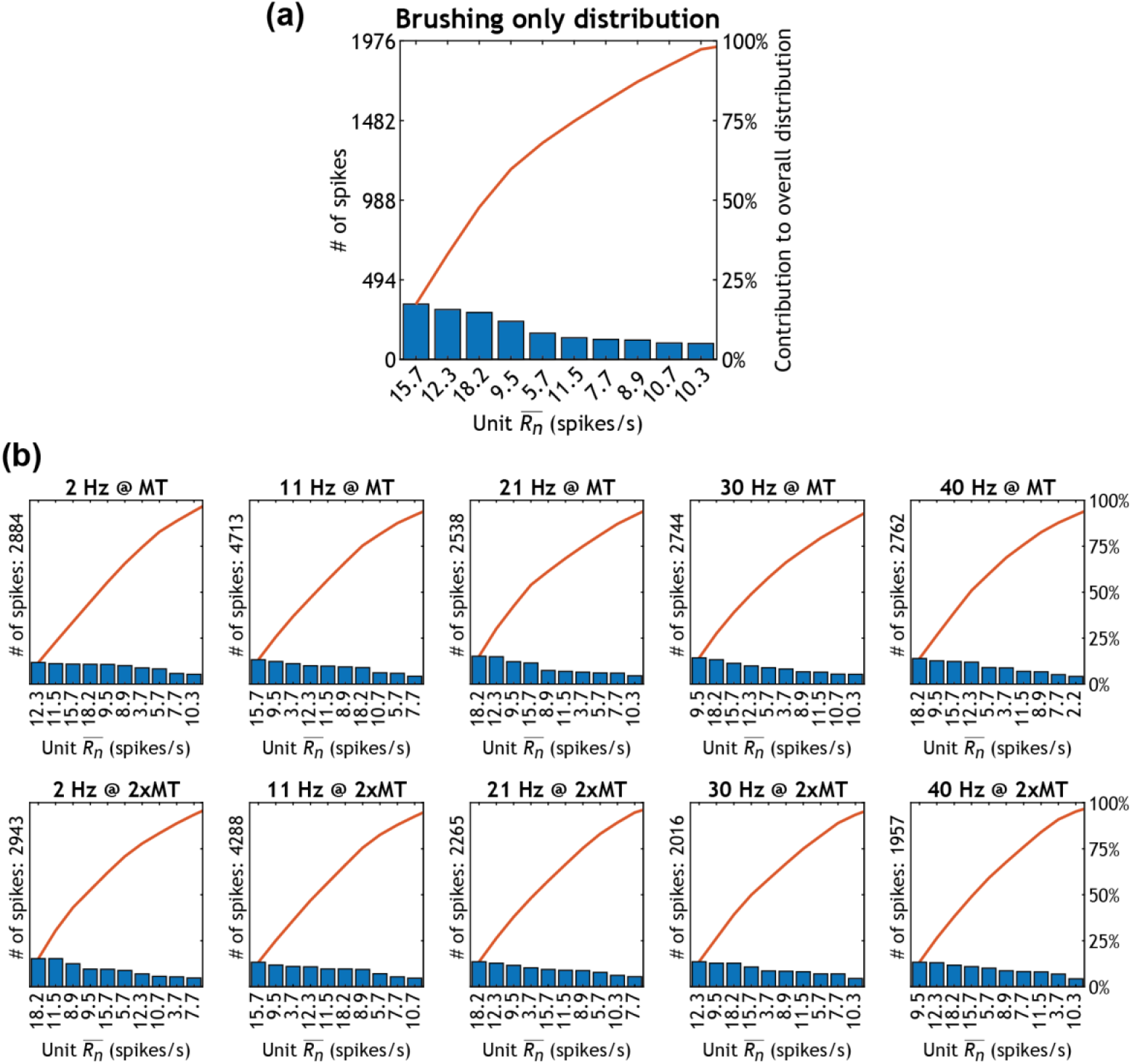
Pareto chart for the data collection session with Animal 1. Bars are labeled by the respective unit’s 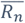. Combined ISIs for the brushing only trial is shown in (a). The total number of spikes making up each electrical stimulation trial’s distribution are listed on the y-axis of plots in (b).

*Figure 6* shows a Pareto chart for all brushing units from the data collection session with Animal 1. Units with 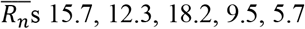, and 11.5 spikes/s were the top 6 contributors to the baseline distribution, and they remained within the top 10 contributors to the combined distribution across all stimulation settings. Likewise, in the first data collection session with Animal 2, units with 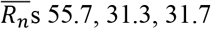, and 30.8 spikes/s made up the top 4 contributors to the baseline distribution, and they remained within the top 10 contributors to the combined distribution across all stimulation settings (*Figure S4*). Due to the different responses of units to each stimulation frequency and amplitude level, the changes between combined *T*_*n*_ distributions and a given electrical stimulation trial’s combined *T*_*e*_ distribution were dependent on individual units’ changes in output. For example, during Animal 1’s data collection session, changes in the ISI distribution of the unit with a 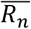 of 12.3 spikes/s during trials that applied 21, 30, and 40 Hz electrical stimulation at 2xMT made this unit one of the top five contributors to both increases and decreases in ISI bin heights in the combined ISI distribution for those trials (*Figure S3*). As an additional example, a unit with a 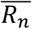 of 2.2 spikes/s, despite not being within the top five contributors to the combined ISI distribution for any stimulation setting, was one of the top five contributors to increases in ISI bin heights in the combined ISI distribution during both 30 Hz stimulation trials. In the data collection session with Animal 1, four units (30%) were within the top five contributors to *change* in at least one trial’s combined ISI distribution despite not being within the top five contributors to any trial’s combined ISI distribution. There were six units (31.6%) from the first Animal 2 data collection session that also shared this property. Only one unit collected from Animal 3 responded to cutaneous brushing and discernably responded directly to electrical stimulation, so Pareto charts for this animal’s data are not shown.

### Individual neural unit responses

Underlying the combined ISI distributions, data from each individual neural unit displayed a wide range of changes in firing behavior in response to electrical stimulation. To quantify changes in output for individual units, we generated two separate yet related log-linear mixed-effect models. We used a given unit’s 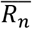, applied electrical stimulation frequency, applied electrical stimulation amplitude level, and the unit’s conduction velocity as predictors for the unit’s response during a trial. We quantified a unit’s response with 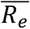 for the first model (*Figure 7b* and *7c*) and change in 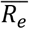 as a percentage of 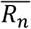 for the second model (*Figure 8b* and *8c*). Between the two models, all fixed effects coefficients were identical except for the coefficient *β*_*n*_ (unit 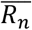), which was 0.582 for the first model (*p*-value, 5.9e-20; 95% confidence interval (CI), [0.462, 0.702]) and −0.418 for the second model (*p*-value, 2.1e-11; 95% CI, [−0.538, −0.298]). While electrical stimulation frequency (*β*_*s*_, −0.054; *p*-value, 5.3e-9; 95% CI, [−0.072, −0.036]) and electrical stimulation amplitude (*β*_*a*_, −0.097; *p*-value, 2.7e-5; 95% CI, [−0.143, −0.052]) were inversely correlated with both metrics of unit response, conduction velocity was not found to have a significant impact (*β*_*v*_, 0.007; *p*-value, 0.90). The effect sizes that were calculated with normalized predictor variable data for the 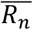 coefficients in each model (0.414 and −0.297, respectively) were over five and three times greater in magnitude, respectively, than the coefficient effect size for electrical stimulation frequency (−0.077). The 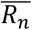 coefficients’ effect sizes were over six and four times greater in magnitude, respectively, than the electrical stimulation amplitude coefficient’s effect size (−0.069). The Linear Mixed-Effects Models section of the Supplemental Materials contains tables for each model and provides the goodness of fit and residuals (*Table S3, Table S4*, and *Figure S7*).

**Figure 7:**
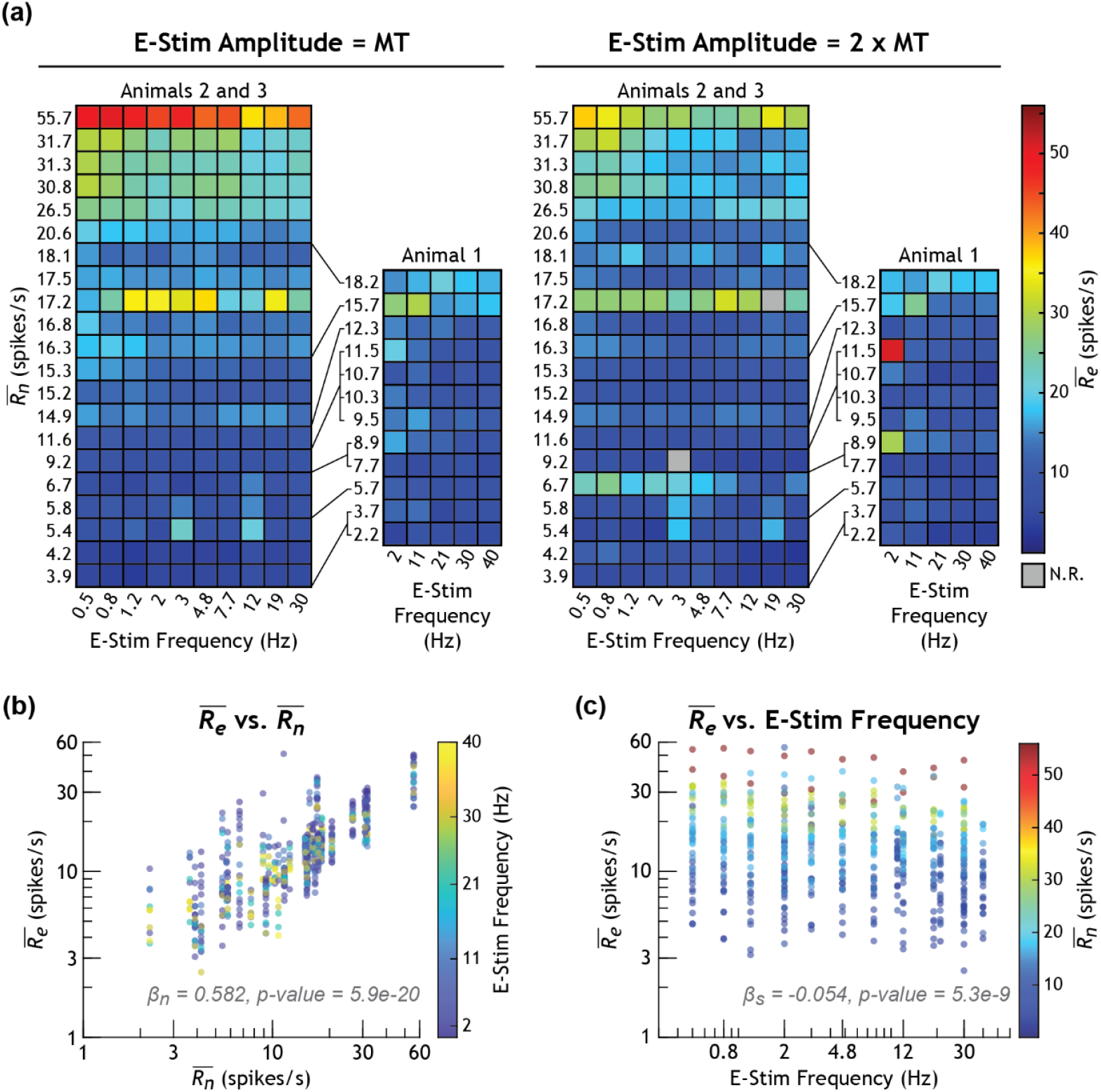
The mean firing rate 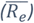 for each unit during each electrical stimulation trial, calculated using the log average of instantaneous firing rates, as a function of each unit’s 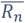 and electrical stimulation frequency. (a) Heatmaps of each unit’s response during each trial. Each row represents the responses of a single unit, where units are ordered from bottom to top by increasing mean underlying firing rate 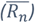. Each unit’s mean underlying firing rate is used as the row label along the left of each heatmap. Each square in the heatmaps represents the response of that unit during a stimulation trial, where the stimulation frequency used for that trial is listed at the bottom of each column. Because the frequencies used in Animals 2 and 3 were different from those used for Animal 1, their unit responses are plotted in separate heatmaps for each stimulation amplitude. The relative order positions of Animal 1’s units within Animal 2 and 3’s units are indicated by the black lines connecting their row labels to the larger heatmaps. (b) Units’ 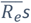 across all stimulation trials plotted against their 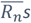. Each dot represents a unit’s response during a trial, and the color of the dot corresponds with the stimulation frequency used for that trial. (c) Units’ mean endpoint firing rates across all stimulation trials plotted against the stimulation frequency used for that trial. Each dot represents a unit’s response during a trial, and the color of the dot corresponds with the underlying firing rate for that unit. The coefficients for each predictor variable in (b) and (c) from the log-linear mixed effects model is included in the plot, along with their p-values. N.R. = No Response.

**Figure 8:**
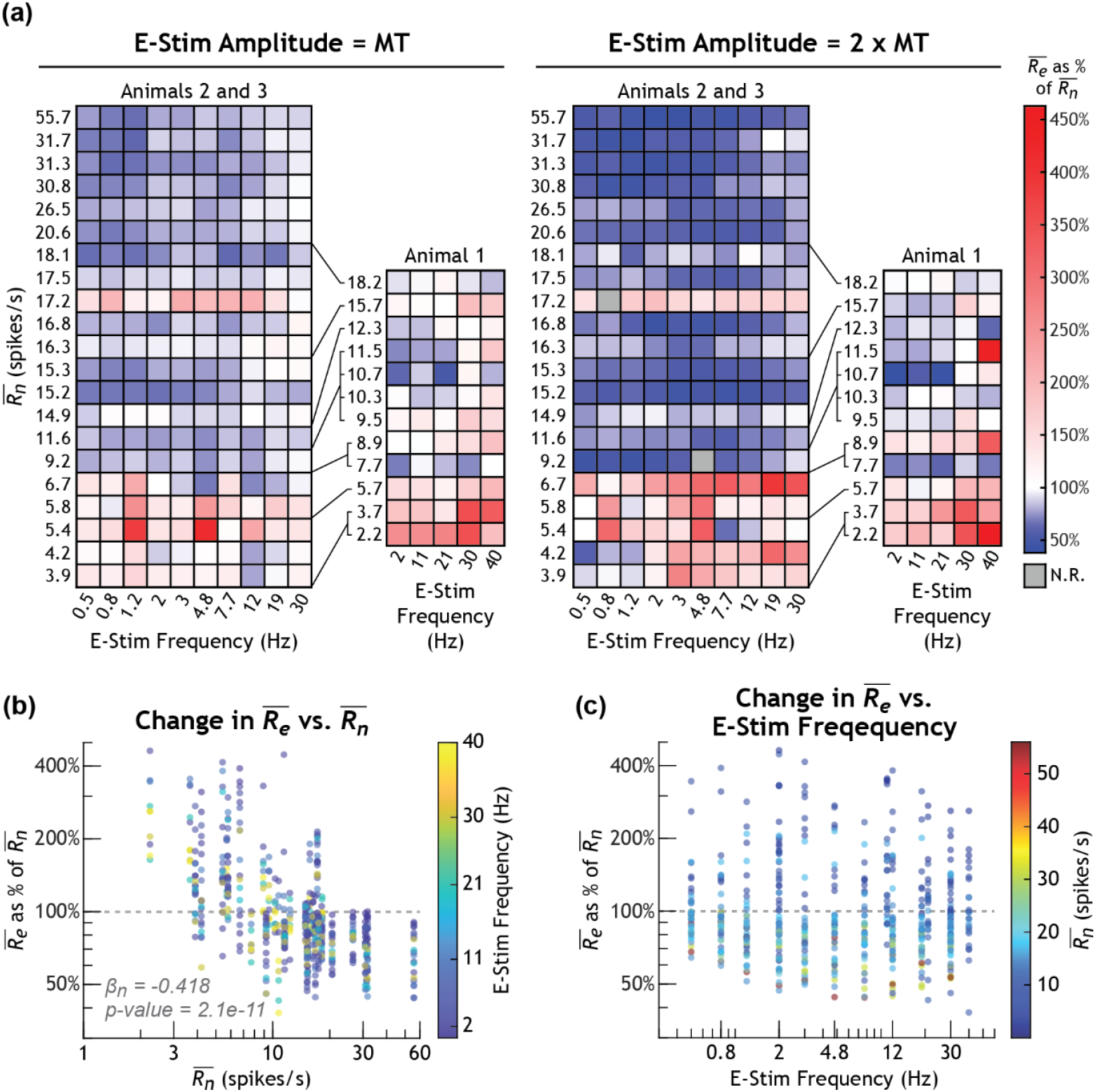
Heatmaps showing each unit’s firing rate during electrical stimulation trials as a percentage of their baseline firing rate. (a) See caption for Figure 6. Here, the color of each square corresponds to the percent change, for that trial, in mean endpoint firing rate 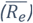 from that unit’s mean underlying firing rate 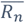. (b) and (c) See caption for Figure 6. Here, each variable is plotted against percent change in 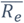. The coefficient for 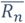 in (b) from the log-linear mixed effects model is included in the plot, along with the p-value. The coefficient and p-value for stimulation frequency (c) was the same as shown in Figure 6. N.R. = No Response.

We quantified how a unit’s 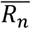 value and the applied electrical stimulation settings influenced its 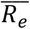 value during trials that applied electrical stimulation (*Figure 7, Figure 8*). At MT amplitude, 14 of the 33 analyzed units (42.4%) increased in their 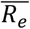 for at least half of the stimulation frequencies tested on a given unit (mean of increasing units’ 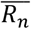 values, 8.3 ± 4.9 spikes/s; range of increasing units’ 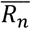 values, 2.2 to 17.2 spikes/s). At 2xMT, this was the case for 10 (30.3%) of the analyzed units (mean of increasing units’ 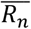 values, 6.4 ± 4.2 spikes/s; range of increasing units’ 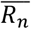 values, 2.2 to 17.2 spikes/s). Alternatively, at MT, the remaining 19 of the 33 units (57.6%) decreased their firing rates for over half of the stimulation frequencies tested on a given unit (mean of decreasing units’ 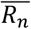 values, 19.7 ± 11.5 spikes/s; range of decreasing units’ 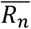 values, 7.7 to 55.7 spikes/s). At 2xMT, this was the case for the remaining 23 units (69.7%) (mean of decreasing units’ 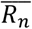 values, 18.6 ± 10.8 spikes/s; range of decreasing units’ 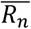 values, 7.7 to 55.7 spikes/s). These increasing and decreasing changes in mean firing rate compared to brushing only trials were more pronounced during 2xMT trials than MT trials (*Figure 8a*).

Interestingly, several units with medium to larger 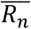 values increased their 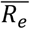 values more prominently during trials using 2xMT amplitude and electrical stimulation frequencies of 1.2 to 12 Hz (units with 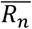 14.9, 15.2, 15.3, 16.3, 16.8, 17.5, 20.6, and 26.5 spikes/s in *Figure 8a*).

We did not observe a significant pattern in the geometric standard deviations of unit *R*_*e*_ distributions across unit 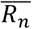 values during MT trials. However, geometric standard deviations of unit *R*_*e*_ distributions increased during 65.6% of unit–trial pairs at MT and 75.9% of unit-trial pairs at 2xMT (*Figure S5*). Many units with larger 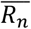 values (16.3 spikes/s and greater in *Figure S5*) displayed a larger increase in standard deviation values for stimulation frequencies 0.8 through 2 Hz during the 2xMT amplitude trials compared to other stimulation frequencies or MT trials. This pattern was also reflected in unit *R*_*e*_ coefficients of variation (which we calculated with the equation for log-normal data, 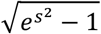, where *s* is the standard deviation of the log-transformed response distribution) across stimulation trials (*Figure S6*).

*Figure 9* shows the analyzed units’ responses for each trial as a function of electrical stimulation amplitude level and conduction velocity. The 2xMT stimulation amplitude (as opposed to MT stimulation amplitude) lead to more prominent increasing and decreasing shifts in 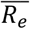 from 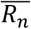.

**Figure 9:**
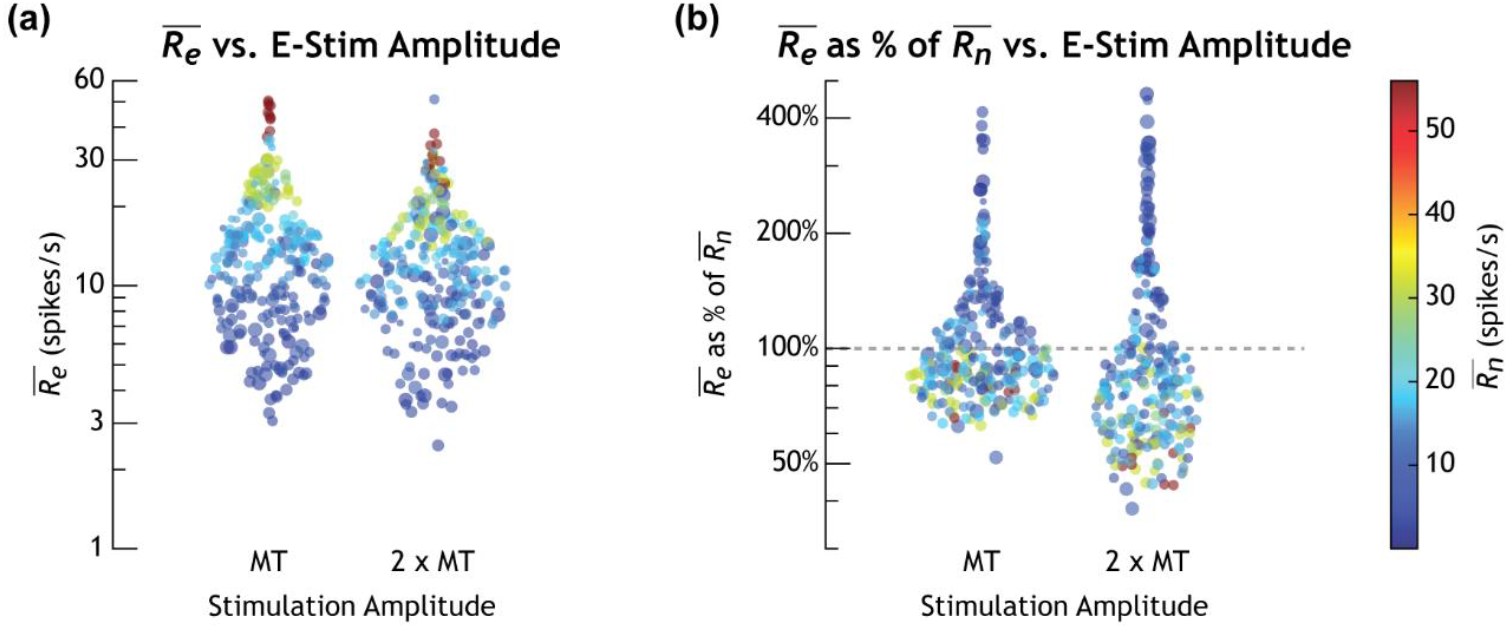
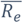 (a) and 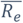 as a percentage of 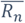 (b) for each unit during each trial plotted against that unit’s electrical stimulation amplitude level. The color of the dot corresponds with the respective unit’s 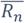 with the color bar on the right. The size of the dot corresponds with unit conduction velocity, where a larger dot represents the response of a unit with a larger conduction velocity.

## Discussion

Peripheral neuromodulation therapies often involve applying electrical stimulation to nerves that have ongoing physiological activity propagating through them. The relationship between electrical stimulation settings and endogenous firing behavior is not well understood, as interactions between ESAPs and PAPs can influence neural output. We quantified the impact of electrical stimulation frequency and amplitude on the output of active peripheral neurons in an acute *in vivo* feline model. Our results demonstrated how both the underlying activity within peripheral nerves and the electrical stimulation settings applied to them can affect the degree and relative increase or decrease in neural output.

### Units collectively respond both in accordance with and against computational predictions

We combined the ISI distributions of all brushing-modulated and electrically stimulated neural units into histograms for each stimulation paradigm to analyze how electrical stimulation changed the overall response of these units. In computational work by Crago and Makowski 2014, their results indicated that, due to summation, the probability of short ISIs 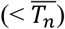 increased when the electrical stimulation frequency was less than that of the mean neural firing rate (Crago and Makowski, 2014). Our experimental results corroborated these computational predictions: a low stimulation frequency — specifically, 2 Hz — resulted in an increase in the probability of very short ISIs (< 0.01 s), which we interpreted as endpoint firing rate summation (*Figure 5*). While this electrical stimulation frequency broadly conserved the propagation of PAPs, the small ISI value range 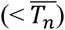 of the distribution was more sensitive to the additive effect of inter-PAP ESAPs.

Conversely, the probability of ISIs that correspond with this (relatively) large *T*_*s*_ value (0.5 s) also increased. Summation, in turn, caused drastic transformations to the firing rate distributions of these units. This can also be seen in the greater increase in the standard deviations and coefficient of variations of *R*_*e*_s from more quickly firing units during trials that applied 2xMT stimulation at 1.2 to 3 Hz (*Figure S5b* and *Figure S6b*). During trials where we applied electrical stimulation frequencies below 2 Hz, because there were fewer ESAPs relative to the number of PAPs, the impact of electrical stimulation on the overall endpoint ISI distribution was minimal during these trials (*Figure S2*).

During trials with the middle-range electrical stimulation frequencies we tested (above 2 Hz and below 30 Hz), the height of ISI probability bins that corresponded with *T*_*s*_ increased, particularly for trials applying 2xMT (*Figures S2* and *S3*). These increased ISI probabilities were likely due to collision block and/or resetting, where the antidromic ESAPs either annihilate PAPs or prevent PAPs from occurring when the antidromic ESAPs reach the sensory end of the neurons. While this was anticipated, there were also increases in ISI bins that corresponded with multiples of *T*_*s*_. This indicates that some units, while they did respond to electrical stimulation, did not respond to every pulse applied. We suspect this may have been due, at least in part, to breathing motion artifacts or vascular pulsation momentarily either shifting the stimulating electrode’s position or applying mechanical stress to the axons (Duncan et al., 2020; Sridharan et al., 2021). These subsequent brief shifts in neural activation should be expected in both *in vivo* animal studies and clinical settings.

Interestingly, the highest stimulation frequencies we applied — particularly 30 and 40 Hz, which were above the means of the combined *R*_*n*_ — showed virtually no change in the probability of ISIs near *T*_*s*_, rather than a substantial increase as computationally predicted in Crago and Makowski 2014 (*Figure 5, S2*, and *S3*). We believe this was likely due to a combination of stimulation entrainment and an increased occurrence of refractory block. Orthodromic ESAPs likely were reaching the endpoint at approximately the same rate as PAPs, while corresponding antidromic ESAPs annihilated PAPs. Additionally, due to entrainment, the likelihood of a PAP passing the site of applied electrical stimulation at the time of an applied pulse increased. This, in turn, would have led to more instances of refractory block as percentage of applied stimulation pulses. With fewer ESAPs reaching the endpoint, electrical stimulation may have had less than the anticipated impact on the overall firing rate distribution at these frequencies.

Lastly, ISIs increased (*R*_*e*_s decreased) throughout the entire distribution during trials where we applied 30 or 40 Hz of electrical stimulation (*Figure 5, S2*, and *S3*). We suspect this was due in part to the high rate of antidromic ESAPs annihilating incoming PAPs, thus eliminating many short ISIs 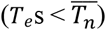. The increase in probability of larger ISIs 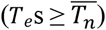 could possibly be due to neural resetting by electrical stimulation or ephaptic effects, where action potentials in separate but adjacent electrically stimulated axons traveled together and slowed for a brief period during their propagation towards the DRG (Capllonch-Juan and Sepulveda, 2020; Katz and Schmitt, 1940). This broad shift in the overall ISI distribution towards larger ISI values during these trials was subtle but consistent between animals.

### Units individually respond in diverse ways depending on their underlying activity

Pareto charts revealed that a select few neural units primarily encoded the physiological neural responses to brushing (*Figure 6*). However, we found changes in which units contributed to recorded responses during the electrical stimulation settings we applied. Further, we identified units that were key contributors to change between brushing only and select electrical stimulation response distributions, despite these units not being primary contributors to the overall responses. This indicated electrical stimulation settings differentially impacted the units encoding neural output. The log-linear mixed effects models constructed from this data revealed the significant impact of underlying physiological activity on both the endpoint firing rate of individual neurons (*Figure 7*) and whether their firing rate increased or decreased (*Figure 8*). Intuitively, the slower firing units within this dataset generally increased their endpoint firing rate in response to electrical stimulation, with a greater proportion of their endpoint action potentials likely being orthodromic ESAPs. The quicker firing units in this dataset decreased their endpoint firing rate in response to stimulation, likely because antidromic ESAPs annihilated a greater proportion of their PAPs. Because of the length of these axons — and thus more time it takes for PAPs to reach the site of electrical stimulation — the endpoint firing rate of these units were effectively capped by high electrical stimulation frequencies. Further, previous studies have demonstrated how antidromic action potentials in primary sensory afferents can delay spikes and reduce the firing frequency of PAPs (Cattaert and Bévengut, 2002; Gossard et al., 1999). The number of antidromic ESAPs that reach the sensory ending and can elicit this behavior depends on both the distance between the stimulating electrode and the ending, in addition to the relative frequencies of neural firing and applied electrical stimulation. Thus, neural resetting by high electrical stimulation frequencies likely also played a role in the reduction of unit firing rates.

The impact of the applied electrical stimulation frequency on a given unit’s response was smaller than the impact of that unit’s underlying firing rate. However, this smaller impact was statistically significant and, surprisingly, negative (*Figures 7 and 8*). While we assumed a linear relationship between the log-transformed rates (of firing activity and applied electrical stimulation), the response residuals did not have a constant variance across the models’ fitted values. Residual variance decreased across fitted values for log-transformed 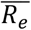 values (*Figure S7c*) and increased across fitted values for the log ratio of 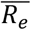 and 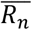 (*Figure S7d*). In other words, model fit was worse for unit responses that had smaller 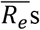 and, correspondingly, worse for unit responses that had larger, positive changes in 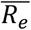 relative to 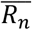. While this is likely due in part to having fewer ISI datapoints for slower firing units, these relationships are also likely non-linear and therefore much more difficult to resolve across different applied electrical stimulation frequencies *in vivo*.

This non-linearity was predicted by Crago and Mackowski 2014, where they described non-monotonic relationships between the ratio of stimulation frequency to 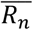 and 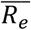. They identified ranges across this latter ratio for which 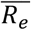, relative to 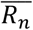, increased or decreased. These relationships depended on additional factors such as inter-site conduction time (e.g., the time it takes a PAP to leave the sensory ending and reach the site of electrical stimulation) and the length of the refractory period for a given axon. Here, we considered the impact of inter-site conduction time in our statistical models by proxy of conduction velocity. However, the log-linear mixed effects models indicated this factor was insignificant for units in our dataset.

Even so, we would like to highlight the role of propagation time between a physiologically transitional region (such as a sensory ending, a synapse in the spinal cord, or a neuromuscular junction) and the location of the stimulating device. Formento 2018, who utilized electrical stimulation of the spinal cord to restore locomotion, identified discrepancies between rat and human subjects for whether spinal cord stimulation silences proprioceptive information important for coordinated motor function (Formento et al., 2018). Humans have longer nerves, thus allowing more propagation time for collision block to occur. Additionally, rats have proprioceptive firing rates that span much higher rates per second, which reduces the likelihood each proprioceptive PAP within a spike volley could become blocked by an antidromic ESAP for a given electrical stimulation frequency. They used both clinical and computational experiments to demonstrate the importance of selecting stimulation paradigms with respect to underlying physiological activity in order to preserve off-target information pertinent to clinical goals.

Felines, although larger than rat animal models, still have shorter lumbar and sacral level nerves than humans by a factor of at least half (O’Bichere et al., 2000; Pirro et al., 2009). This means, in humans, it is more likely that quickly firing units will decrease in response to stimulation if the applied frequency is lower than the physiological firing rate. Additionally, low frequency stimulation will lead to summation less often than was observed in this study when the stimulating electrode is farther from the PAP source. The underlying neural firing rate at which the change in mean endpoint firing rate of a neuron switches from positive to negative in response to electrical stimulation was found to be around 9 spikes/s in this study (*Figure 8*). We anticipate this transitionary value would be lower in subjects where the stimulating electrode was placed further from the source of PAPs.

### Clinical implications

Elucidating the mechanisms of peripheral neuromodulation devices might help streamline patient programming and improve device outcomes. The settings used by different applications of peripheral nerve stimulation vary across frequency, pulse width, and device placement, to name a few clinically controllable factors. What configurations are ultimately used by patients varies, but recommended ranges for frequency and pulse width based on extensive clinical research usually provide a starting point.

Overactive bladder (OAB) and underactive bladder (UAB) are generally both treated with an sacral neuromodulation (SNM) stimulation frequency of 14 Hz (Cohn et al., 2017; Thomas and Hashim, 2024). How matching treatment paradigms are often able to treat contrasting pathologies remains unclear. Previous literature speculate that SNM “normalizes” the activity of continence and micturition neural circuits (Gill et al., 2017; Thomas and Hashim, 2024). Our findings may support this hypothesis. It is possible that patients with OAB or UAB disorders might have insufficient or excess neural activity within their micturition neural circuits that SNM normalizes.

Similarly, uncovering the mechanisms of vagus nerve stimulation for its various uses requires considering the relationship between (a) clinically controllable factors such as stimulation frequency and implantation location and (b) underlying neural activity and the desired response of downstream organs or central nervous system pathways. Clinicians and researchers might consider how off-target effects could be mitigated by limiting the range of frequencies to be tested clinically.

Peripheral neuromodulation patients who do not see improvement in their symptoms could potentially benefit from more drastic, rather than minute, adjustments in stimulation frequency. For any peripheral neuromodulation therapy, inconsistent clinical outcomes should be analyzed with respect to potential variability in underlying activity and relative inter-site distances. Clinical implications of animal studies should also be interpreted with respect to these factors, as was done by Formento 2018.

### Limitations and future directions

The methodology for this *in vivo* study could be improved in several ways. Longer trial recordings would have led to better measurements for unit firing rate distributions, particularly for slower firing units. Repeating brushing only trials between and after completing electrical stimulation trials would provide information on the stability of units’ physiological firing behavior across trial sets. Data collection sessions repeated on additional animals would provide more units to be used in the log-linear mixed effects model. Electrical stimulation settings for trials within a data collection session were not randomized and the time between trial recordings was less than one minute. It is possible that DRG cross-depolarization led to unit firing rates recorded during later trials (which applied higher electrical stimulation frequency and amplitude) being moderately inflated compared to those during earlier trials. Future directions for this work could involve mitigating these issues, in addition to repeating this study paradigm on additional peripheral nerves, with additional forms of physiological stimuli and with additional stimulation parameters.

### Conclusions

We sought to quantify how underlying neural firing activity within peripheral sensory axons can impact targeted neural output during electrical stimulation. This was the first study to quantify these neural unit-level relationships in an *in vivo* setting. Log-linear mixed effects models revealed that for individual units, underlying firing rates had the most significant impact on both mean endpoint firing rate and whether firing rates increased or decreased. Compared to trials where action potentials were only evoked by cutaneous brushing, the mean endpoint firing rates of slower spiking units generally increased in response to electrical stimulation, and those of quicker spiking units generally decreased. These findings provide translational context for peripheral neuromodulation research. We highlighted how ongoing neurophysiological activity — whether pathological or off-target — interacts with responses evoked by electrical stimulation and thus can impact clinical outcomes.

## Acknowledgements

We would like to thank Chris Andrews for his extensive guidance on the statistical methods used in this study. We also thank the Unit for Laboratory Animal Medicine at the University of Michigan for their services. Lauren R. Madden was funded by the J. Robert Beyster Computational Innovation Graduate Fellows Program and by a Graduate Assistance in Areas of National Need (GAANN) Fellowship from the Department of Education (P200A220004). This study was also funded by a National Science Foundation (NSF) Career Award 1653080. Finally, we would like to acknowledge Pat Crago, who inspired this study in a conversation with Tim Bruns.

