## Supplemental for "Physiological activity within peripheral nerves influences neural output in response to electrical stimulation: an *in vivo* study"

### Supplemental Materials

*Table S1: Animal data collection sessions*

| Animal | Data collection session | Side and DRG | Trial set # | Cutaneous brushing region | # and range of e-stim freqs. | Pulse duration |
|---|---|---|---|---|---|---|
| Animal 1 | Session 1 | Right S1 and S2 | 1 | Anal | 5 frequencies, 2 - 40 Hz | 200 µs |
| | | | 2 | Scrotal | 5 frequencies, 2 - 40 Hz | 200 µs |
| | | | 3 | Perineal | 5 frequencies, 2 - 40 Hz | 200 µs |
| Animal 2 | Session 1 | Left S1 and S2 | 1 | Perineal | 10 frequencies, 0.5 - 30 Hz | 200 µs |
| | | | 2 | Anal | 10 frequencies, 0.5 - 30 Hz | 200 µs |
| | Session 2 | Left S1 | 1 | Scrotal | 10 frequencies, 0.5 - 30 Hz | 210 µs |
| Animal 3 | Session 1 | Left S1 and S2 | 1 | Perineal | 10 frequencies, 0.5 - 30 Hz | 210 µs |
| | Session 2 | Left S1 and S2 | 1 | Perineal | 10 frequencies, 0.5 - 30 Hz | 210 µs |

*Table S2: Number of units and their respective contribution (indicated by row label) across all trials (indicated by the column header) to the combined ISI distributions*

| | **All settings** | | **No stim & MT trials** | | **No stim & 2xMT trials** | |
|---|---|---|---|---|---|---|
| Contribution ranking | Animal 1 (12 units) | Animal 2 (19 units) | Animal 1 (12 units) | Animal 2 (19 units) | Animal 1 (12 units) | Animal 2 (19 units) |
| **Top 5** | 2 (16.7%) | 3 (15.8%) | 3 (25%) | 3 (15.8%) | 2 (16.7%) | 4 (21.7%) |
| **Bottom 5** | 3 (25%) | 2 (10.5%) | 4 (33.3%) | 2 (10.5%) | 3 (25%) | 3 (15.8%) |

(a)

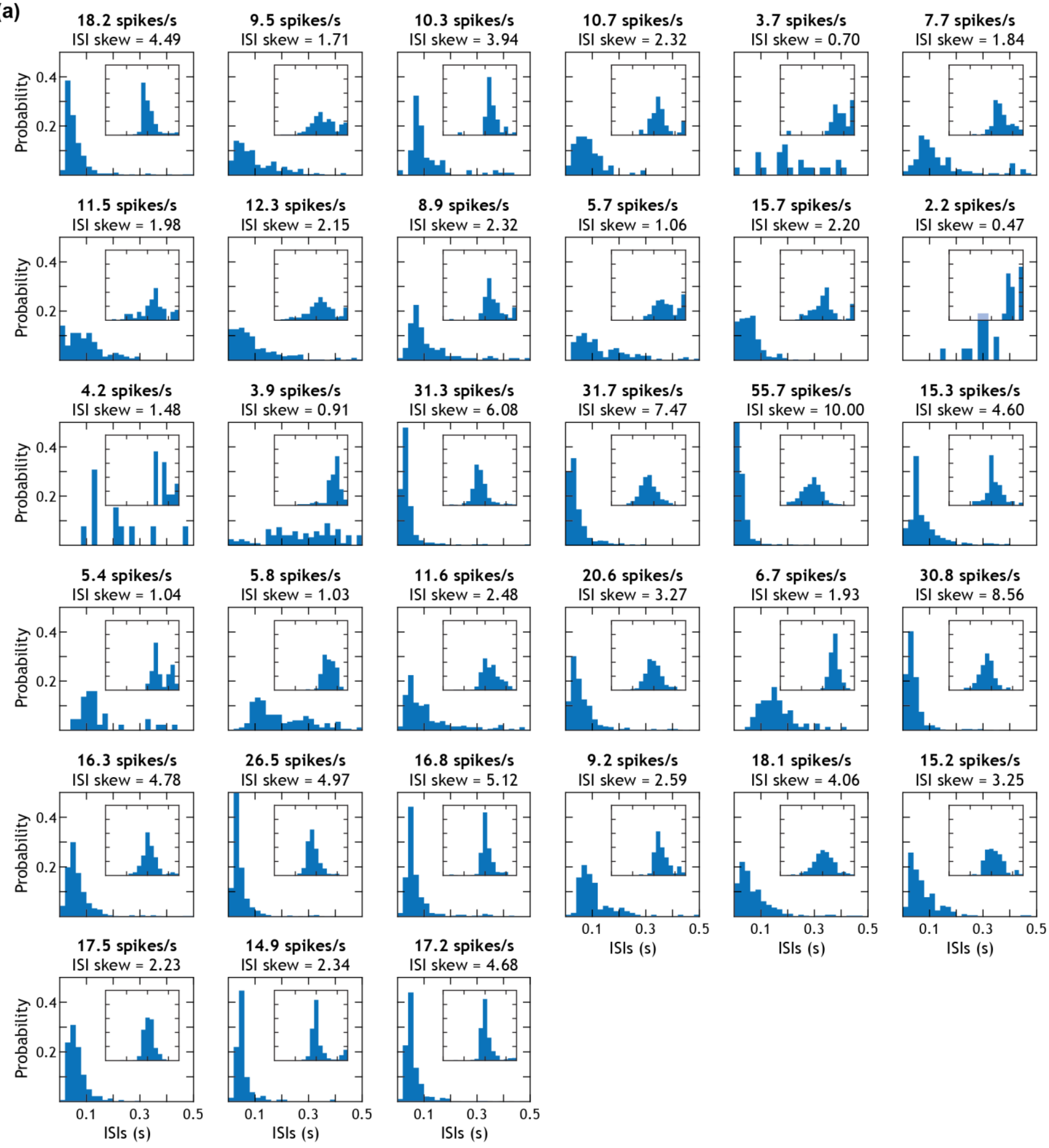

18.2 spikes/s
ISI skew = 4.49
9.5 spikes/s
ISI skew = 1.71
10.3 spikes/s
ISI skew = 3.94
10.7 spikes/s
ISI skew = 2.32
3.7 spikes/s
ISI skew = 0.70
7.7 spikes/s
ISI skew = 1.84
11.5 spikes/s
ISI skew = 1.98
12.3 spikes/s
ISI skew = 2.15
8.9 spikes/s
ISI skew = 2.32
5.7 spikes/s
ISI skew = 1.06
15.7 spikes/s
ISI skew = 2.20
2.2 spikes/s
ISI skew = 0.47
4.2 spikes/s
ISI skew = 1.48
3.9 spikes/s
ISI skew = 0.91
31.3 spikes/s
ISI skew = 6.08
31.7 spikes/s
ISI skew = 7.47
55.7 spikes/s
ISI skew = 10.00
15.3 spikes/s
ISI skew = 4.60
5.4 spikes/s
ISI skew = 1.04
5.8 spikes/s
ISI skew = 1.03
11.6 spikes/s
ISI skew = 2.48
20.6 spikes/s
ISI skew = 3.27
6.7 spikes/s
ISI skew = 1.93
30.8 spikes/s
ISI skew = 8.56
16.3 spikes/s
ISI skew = 4.78
26.5 spikes/s
ISI skew = 4.97
16.8 spikes/s
ISI skew = 5.12
9.2 spikes/s
ISI skew = 2.59
18.1 spikes/s
ISI skew = 4.06
15.2 spikes/s
ISI skew = 3.25
17.5 spikes/s
ISI skew = 2.23
14.9 spikes/s
ISI skew = 2.34
17.2 spikes/s
ISI skew = 4.68
Probability
0.4
0.2
0.1
0.3
0.5
ISIs (s)

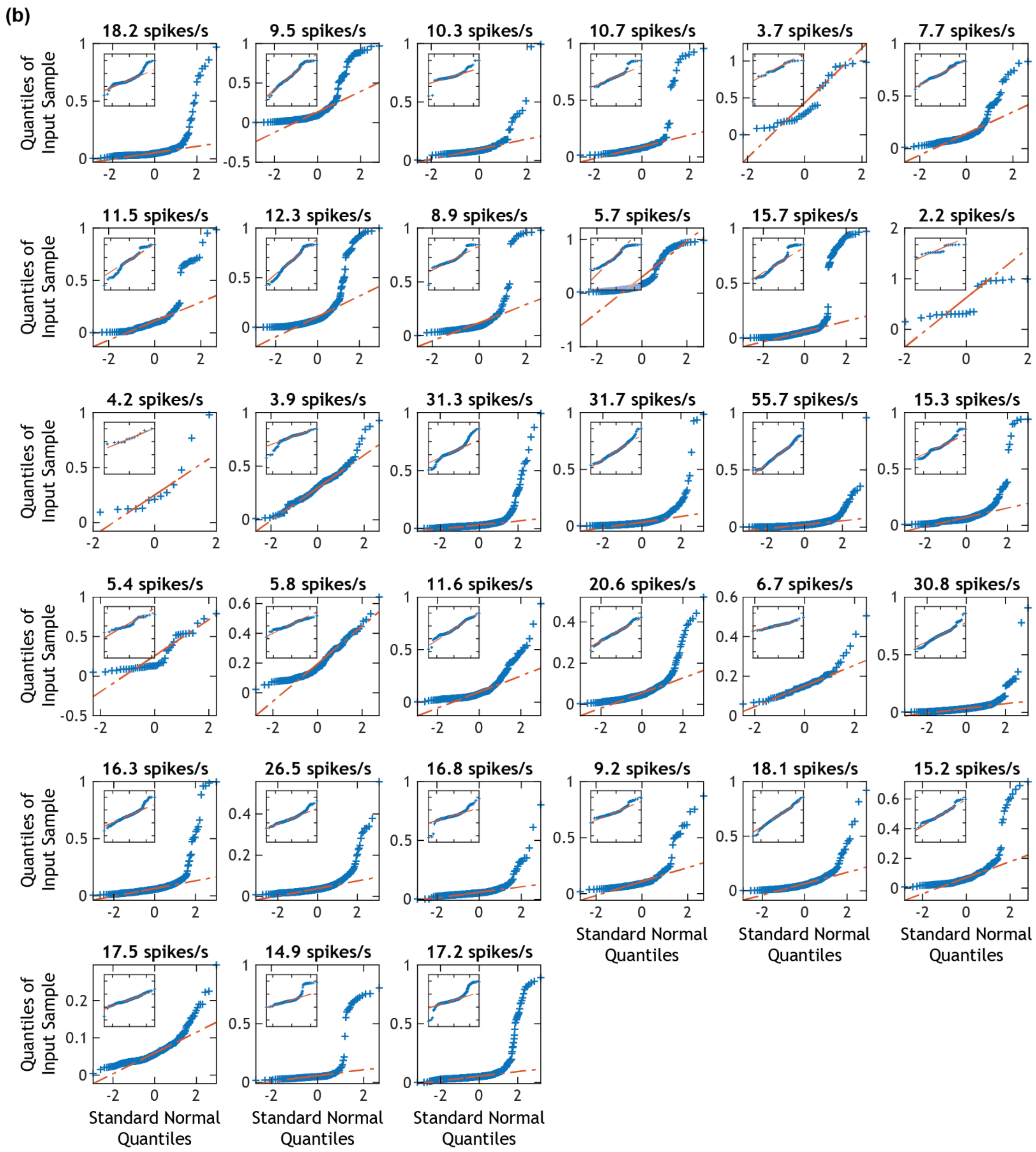


<u>Figure S1:</u> *(a) Brushing-only trial ISI distributions for all 33 brushing-modulated units that were confirmed to be activated by electrical stimulation. The title of each plot gives the unit's* $\overline{\mathrm{R_n}}$ *and the skewness of the ISI (i.e.* $\mathrm{T_n}$ *) distribution. Each individual plot has an inset showing the natural log transformation of the same data. The x-axis for each log plot ranges from -7 to 0, and the y-axis is the same as the main plots. (b) Q-Q plots for the brushing-only trial ISI distributions for all 33 units. Each individual plot has an inset showing the Q-Q plot for the natural log transformation of the same data. The x-axis for each inset plot is the same as the respective main plot, and the y-axis ranges from -7 to 1.*

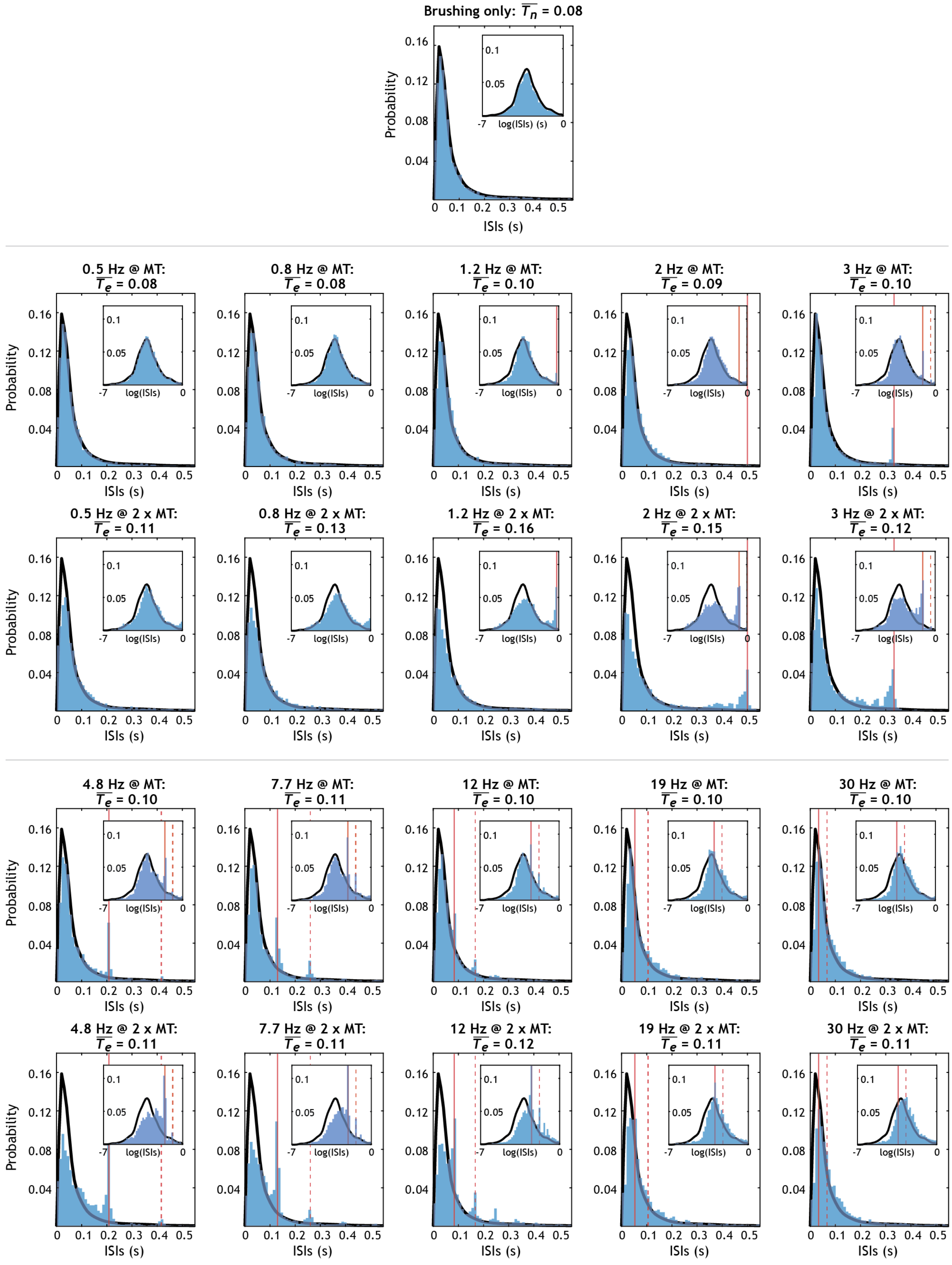


*<u>Figure S2:</u> Combined ISI distributions for all units from the first session with Animal 2 that responded to*

*brushing and electrical stimulation. Titles include e-stim settings and $\overline{T}_e$, the geometric mean, of the distributions. Insets contain the natural log transformation of the same data. Solid red vertical lines denote $T_s$, which is the inverse of electrical stimulation frequency. Dotted red vertical lines denote $2 \times T_s$.*

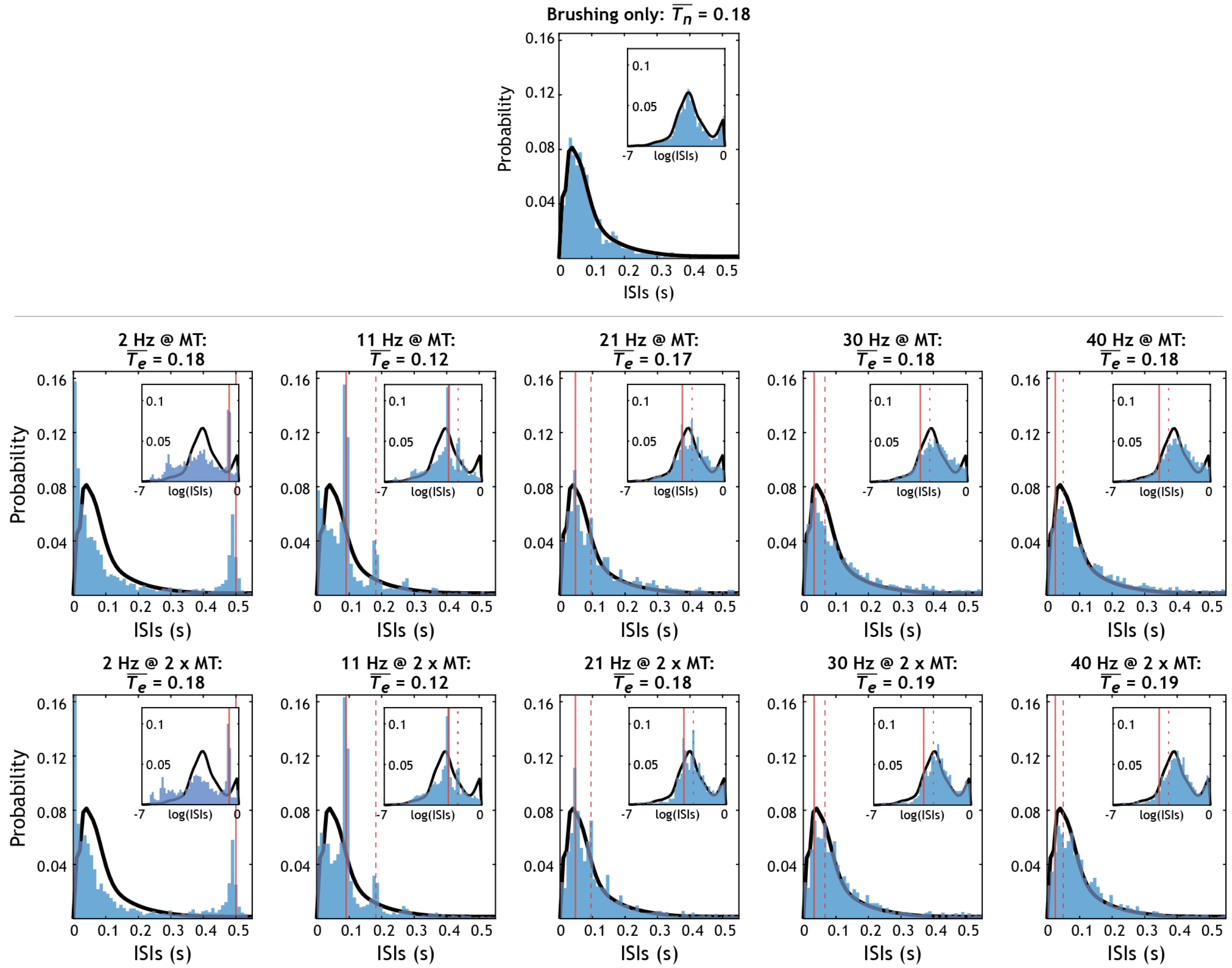


*<u>Figure S3:</u> Combined ISI distributions for all units from the first session with Animal 1 that responded to brushing and electrical stimulation. Titles include e-stim settings and $\overline{T}_e$, the geometric mean, of the distributions. Insets contain the natural log transformation of the same data. Solid red vertical lines denote $T_s$, which is the inverse of electrical stimulation frequency. Dotted red vertical lines denote $2 \times T_s$.*

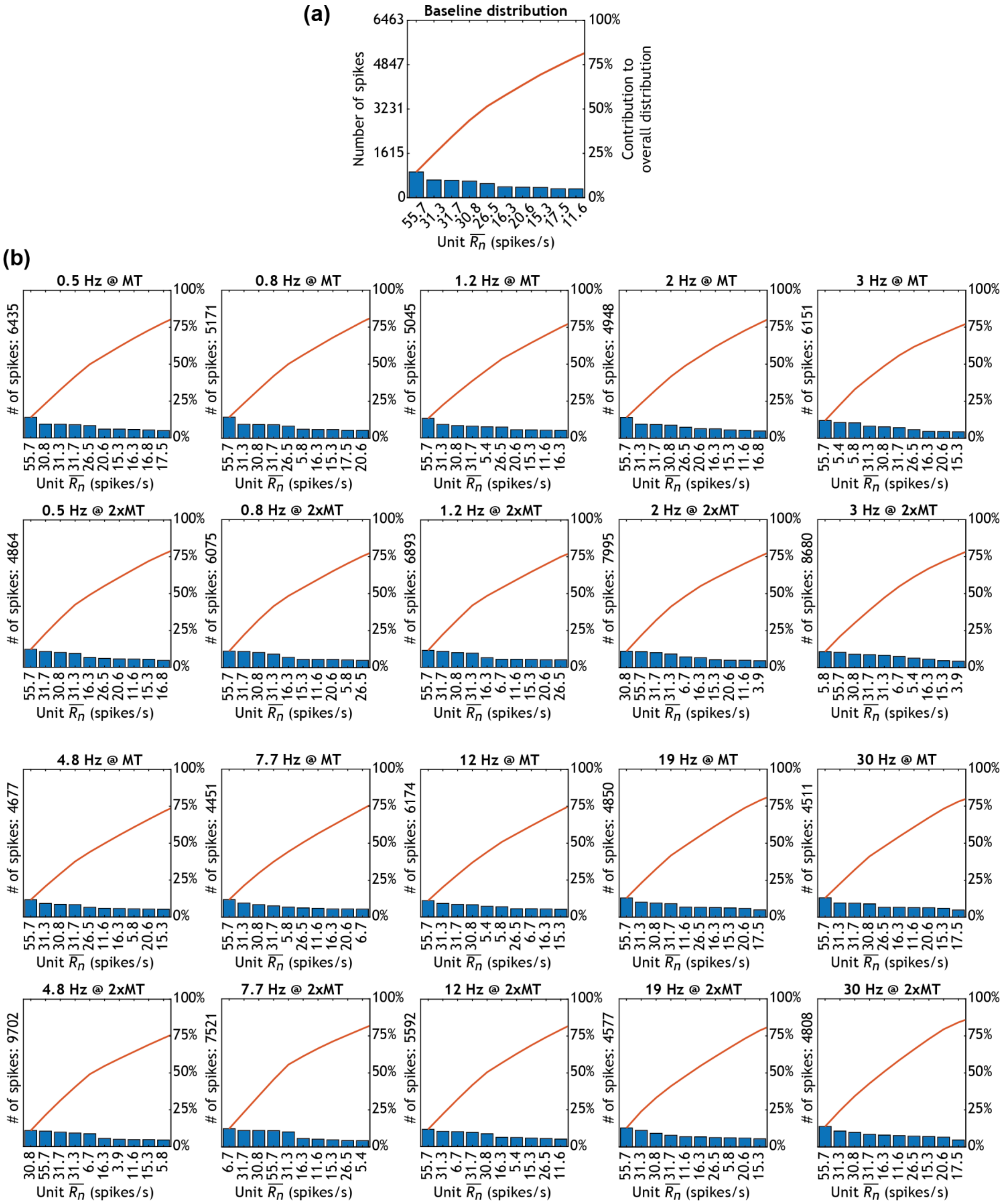

*<u>Figure S4:</u> Pareto chart for the first data collection session with Animal 2, corresponding to Figure 6. Bars are labeled by the respective unit's* $\overline{R_n}$.

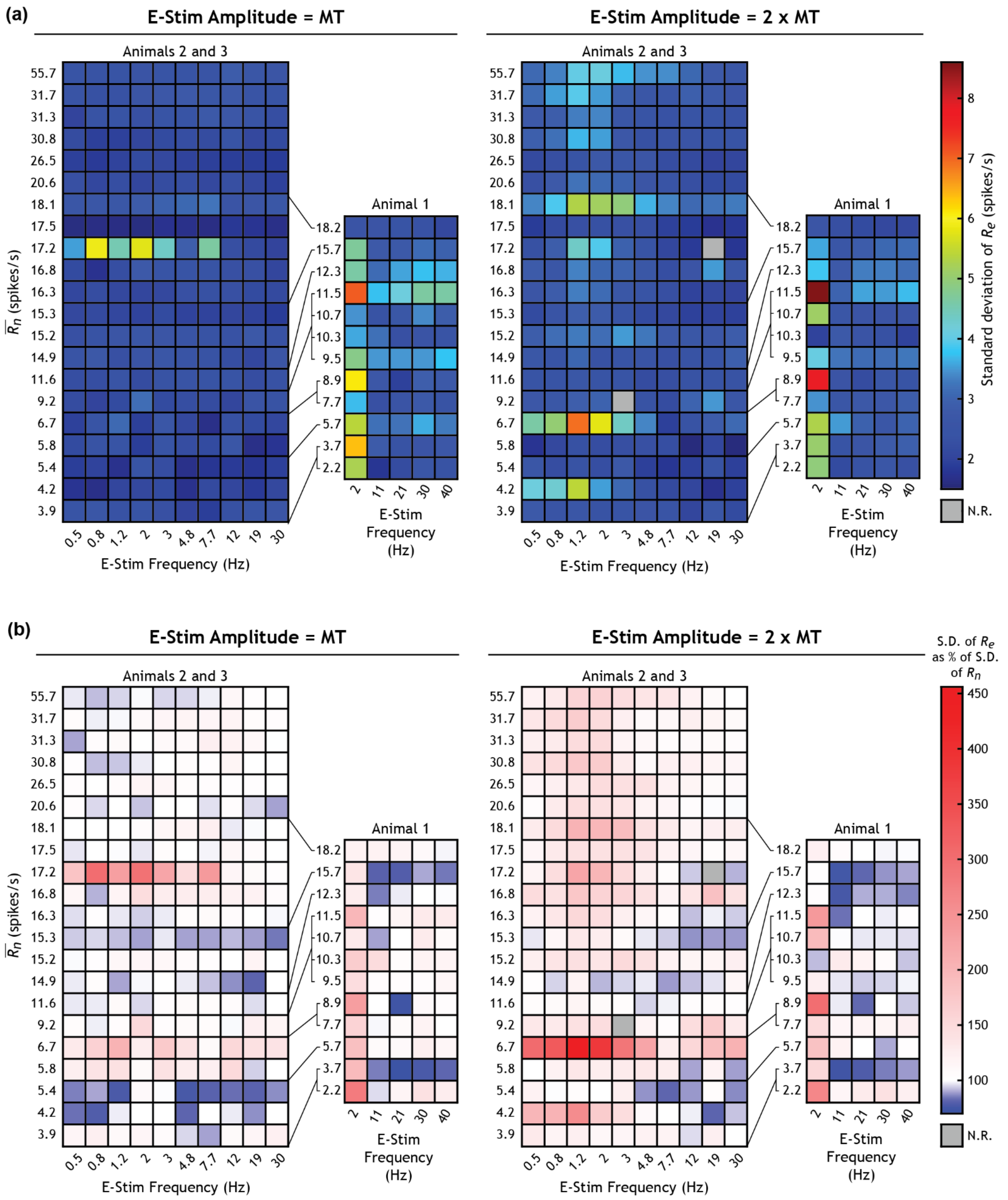


*Figure S5:* Standard deviations (S.D.) of $\mathrm{R_e}$ and changes in the standard deviations of $\mathrm{R_e}$ of all 33 brushing modulated and electrically stimulated units, ordered by $\overline{\mathrm{R_n}}$. (a) The geometric standard deviation of $\mathrm{R_e}$ for each unit during each trial. (b) The geometric standard deviation of $\mathrm{R_e}$ as a percentage of the geometric standard deviation of $\mathrm{R_n}$ calculated via their log ratio/log difference. N.R. = No Response.

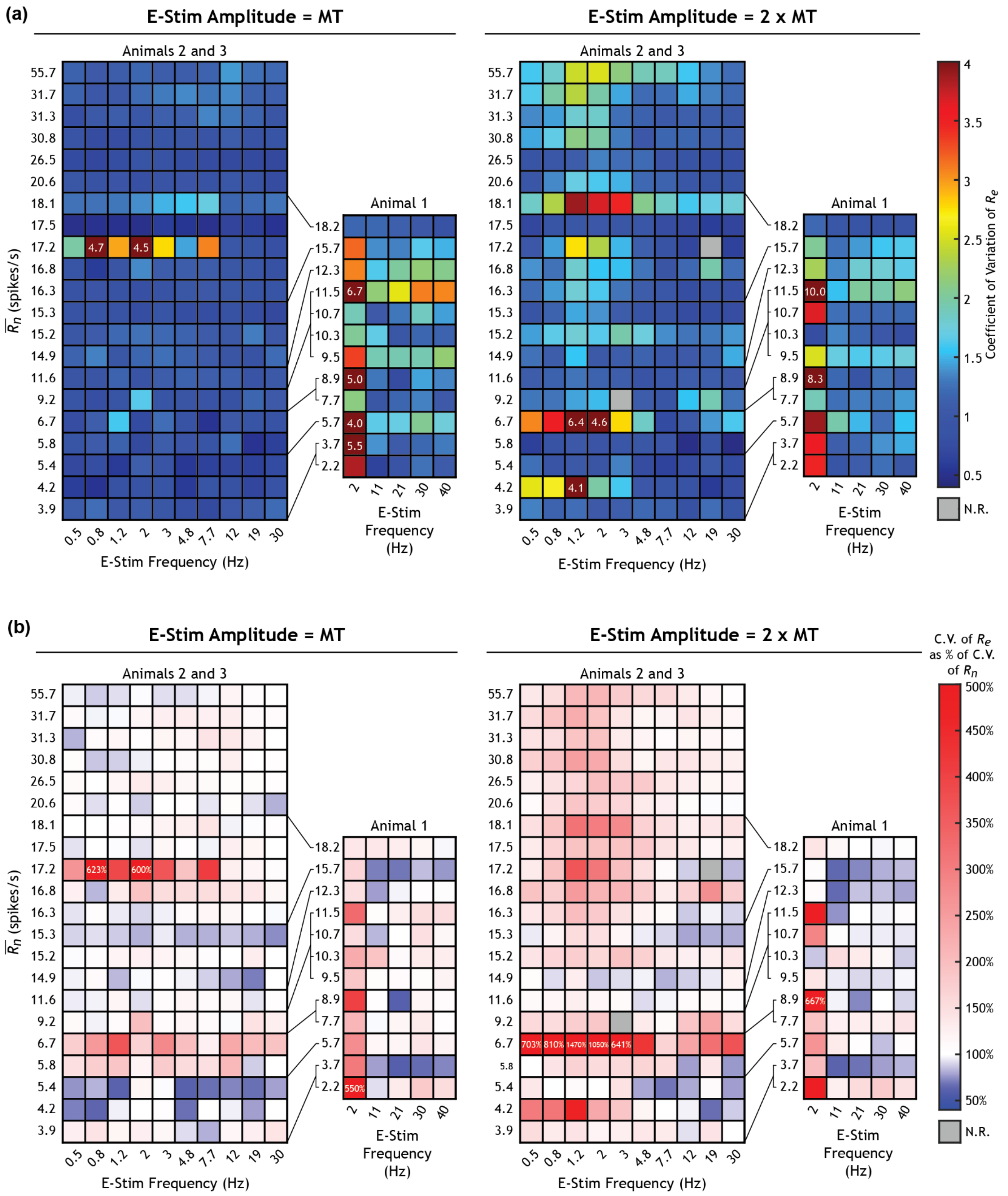


*<u>Figure S6:</u> Coefficients of variation (C.V.) of* $\mathrm{R_e}$ *and changes in the coefficients of variation of* $\mathrm{R_e}$ *of all 33 brushing modulated and electrically stimulated units, ordered by* $\overline{\mathrm{R_n}}$*. (a) The coefficients of variation* $\mathrm{R_e}$ *of for each unit during each trial. (b) The coefficient of variation of* $\mathrm{R_e}$ *as a % of the coefficient of variation of* $\mathrm{R_n}$*. For unit trial responses that exceed the color bar limits, the value is included on the plot. N.R. = No Response.*

**Linear Mixed-Effects Models**

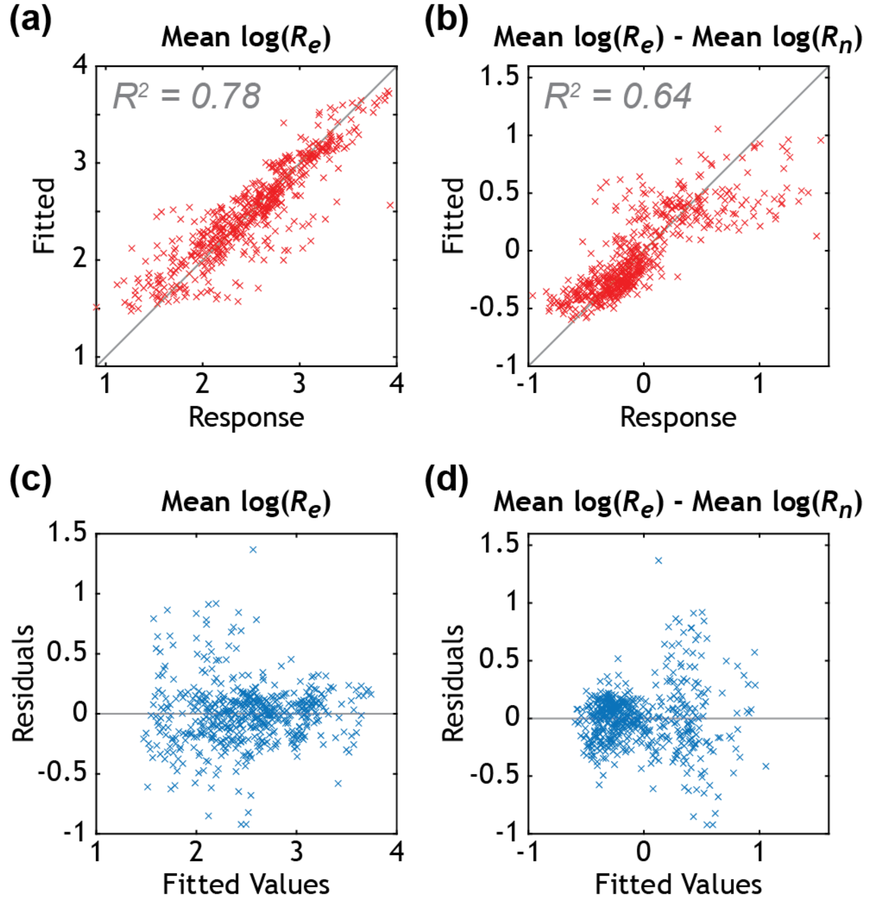


<u>Figure S7:</u> *Plots (a) and (b) show goodness of fit for the first and second linear mixed-effects models, respectively. Plots (c) and (d) show model residuals across fitted values for the first and second models, respectively.*

*Table S3:* First model: Predictors for $\overline{R_e}$ (*Equation 2*)

$$mean[\ln(R_e)] = \beta_0 + \beta_n \,\text{mean}[\ln(R_n)] + \beta_s \ln(R_s) + \beta_a a + \beta_v v + \gamma(1|u) + \varepsilon$$

| **Model Information:** | |
|---|---|
| Number of observations | 538 |
| Fixed effects coefficients | 5 |
| Random effects coefficients | 33 |
| Covariance parameters | 2 |

**Fixed effects coefficients (95% Confidence Intervals):**

| Name | Estimate | Standard Error | Test Statistic | Degrees of Freedom | $p$-value | Lower CI bound | Upper CI bound |
|---|---|---|---|---|---|---|---|
| Intercept ($\beta_0$) | 1.263 | 0.2064 | 6.1186 | 533 | 1.8296e-09 | 0.8575 | 1.6685 |
| $\beta_s$ | -0.0539 | 0.0091 | -5.9351 | 533 | 5.2923e-09 | -0.0717 | -0.0361 |
| $\beta_a$ | -0.0975 | 0.0230 | -4.2325 | 533 | 2.721e-05 | -0.1427 | -0.0522 |
| $\beta_v$ | 0.0072 | 0.0577 | 0.1248 | 533 | 0.9008 | -0.1061 | 0.1205 |
| $\beta_n$ | 0.5818 | 0.0611 | 9.5215 | 533 | 5.9153e-20 | 0.4618 | 0.7019 |

**Random effects covariance parameters (95% Confidence Interval):**

| | Type | Estimate | Lower CI bound | Upper CI bound |
|---|---|---|---|---|
| *Group: Unit* | | | | |
| Intercept | Stnd. dev. | 0.2275 | 0.1746 | 0.2964 |
| *Group: Error* | | | | |
| Residual stnd. dev. | | 0.2670 | 0.2510 | 0.284 |

Ordinary $R^2$: 0.7811
Adjusted $R^2$: 0.7795

*Table S4:* Second model: Predictors for $\overline{R_e}$ as a % of $\overline{R_n}$ (*Equation 3*)

$$mean[\ln(R_e)] - \text{mean}[\ln(R_n)] = \beta_0 + \beta_n \text{mean}[\ln(R_n)] + \beta_s \ln(R_s) + \beta_a a + \beta_v v + \gamma(1|u) + \varepsilon$$

| **Model Information:** | |
|---|---|
| Number of observations | 538 |
| Fixed effects coefficients | 5 |
| Random effects coefficients | 33 |
| Covariance parameters | 2 |

**Fixed effects coefficients (95% Confidence Intervals):**

| Name | Estimate | Standard Error | Test Statistic | Degrees of Freedom | $p$-value | Lower CI bound | Upper CI bound |
|---|---|---|---|---|---|---|---|
| Intercept ($\beta_0$) | 1.263 | 0.2064 | 6.1186 | 533 | 1.8296e-09 | 0.8575 | 1.6685 |
| $\beta_s$ | -0.0539 | 0.0091 | -5.9351 | 533 | 5.2923e-09 | -0.0717 | -0.0361 |
| $\beta_a$ | -0.0975 | 0.0230 | -4.2325 | 533 | 2.721e-05 | -0.1427 | -0.0522 |
| $\beta_v$ | 0.0072 | 0.0577 | 0.1248 | 533 | 0.9008 | -0.1061 | 0.1205 |
| $\beta_n$ | -0.4182 | 0.0611 | -6.8436 | 533 | 2.1293e-11 | -0.5382 | -0.2981 |

**Random effects covariance parameters (95% Confidence Interval):**

| | Type | Estimate | Lower CI bound | Upper CI bound |
|---|---|---|---|---|
| *Group: Unit* | | | | |
| Intercept | Stnd. dev. | 0.2275 | 0.1746 | 0.2964 |
| *Group: Error* | | | | |
| Residual stnd. dev. | | 0.2670 | 0.2510 | 0.284 |

Ordinary $R^2$: 0.6472
Adjusted $R^2$: 0.6446
